# Rational design of disordered proteins for systematic sequence-to-function investigation

**DOI:** 10.1101/2023.10.29.564547

**Authors:** Kara Hunter, Trevor Brandt, Karina Guadalupe, Kavindu Kolamunna, Jeffrey M. Lotthammer, Nora M. Shamoon, Brooke Nicholson, Lea Day, Alec Martinez, Alex S. Holehouse, Shahar Sukenik, Ryan J. Emenecker

**Author notes:** These authors contributed equally to the work (Kara Hunter, Trevor Brandt, Karina Guadalupe).

## Abstract

Despite lacking a stable three-dimensional structure, intrinsically disordered protein regions (IDRs) are ubiquitous across all kingdoms of life and play essential cellular roles. While rational design of folded proteins has seen substantial recent progress, our ability to design IDRs remains limited. Here, we present GOOSE, a comprehensive computational framework for the rational design of IDRs. GOOSE’s versatility and throughput enable us to test thousands of IDR sequences, revealing distinct sequence-ensemble-function relationships. By using GOOSE to rapidly explore these relationships, we design IDRs that can alter their ensembles in response to cell volume changes, create scaffolds for the self-assembly of specific clients, and protect cells from desiccation. Taken together, this work highlights how sequence-function relationships are encoded in IDRs and provides a powerful tool for further exploration.

## INTRODUCTION

Intrinsically disordered proteins and protein regions (IDRs) are found in over 70% of human proteins and play a variety of essential roles^11^. While all proteins exist in an ensemble of conformations, IDRs are defined by the dissimilarity between the 3D configurations of their ensemble, which exists in a dynamic collection of rapidly interconverting and structurally-distinct conformations (**Fig. 1A**)^1–3^. Although “structurally disordered,” IDRs still possess sequence-encoded local intramolecular and long-range intra- and intermolecular interactions that define structural biases within IDR ensembles. Moreover, a growing body of work has demonstrated how such IDR-mediated intra- and intermolecular interactions determine molecular function across various cellular processes^1–3^.

**Figure 1.**
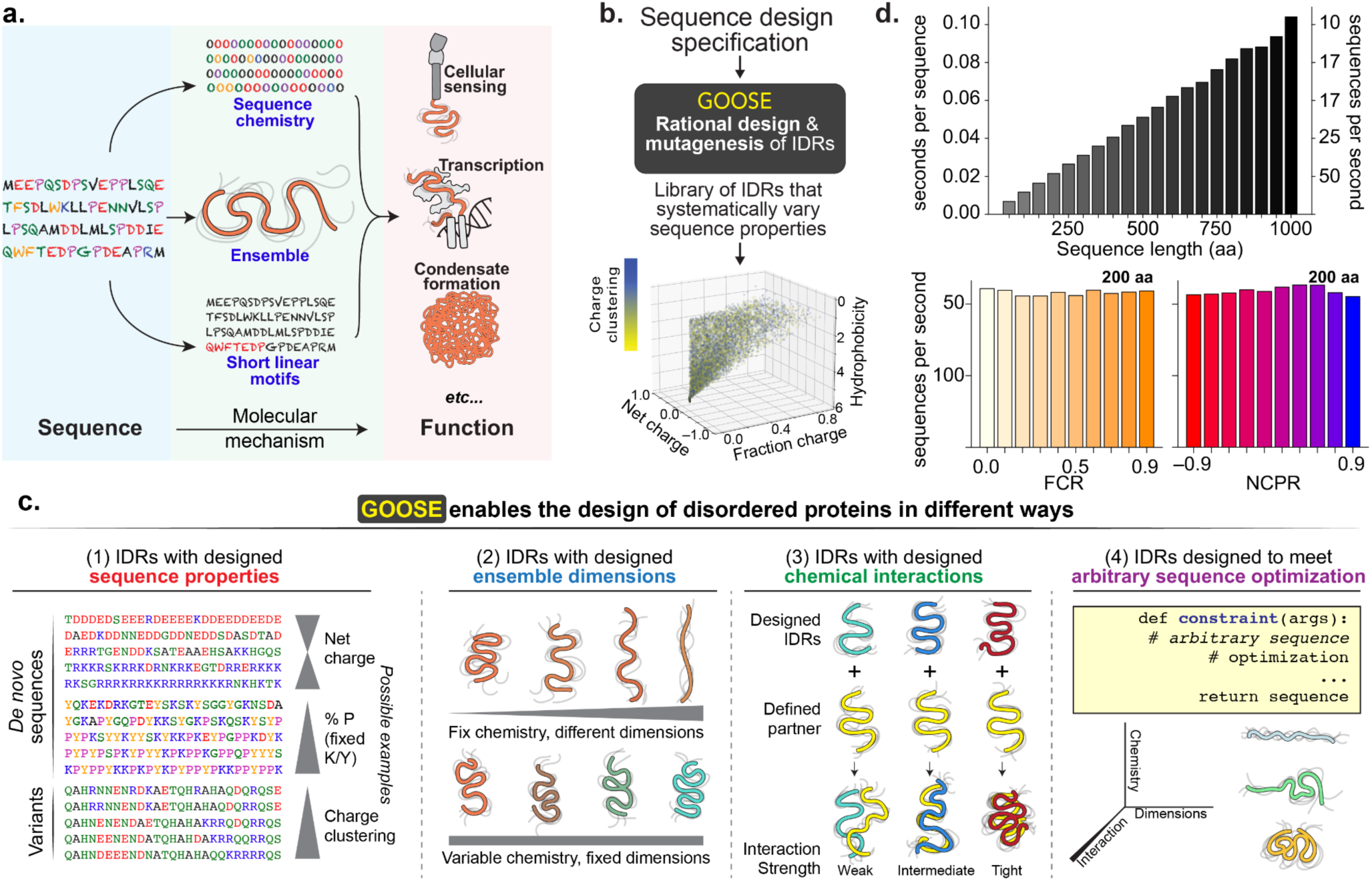
**(A)** IDR function is driven by a combination of sequence-specific effects, ensemble properties, and sequence chemistry. **(B)** GOOSE enables the rational design of disordered proteins that systematically vary the sequence properties of features. **(C)** GOOSE enables the rapid generation of IDR libraries in seconds, enabling high-throughput exploration of sequence-function relationships. GOOSE’s design algorithm enables the generation of thousands of sequences per minute. **(D)** GOOSE enables the design of IDRs with specific sequence properties, ensemble properties, chemically specific intermolecular interactions, or arbitrary design objectives.

Despite their importance, our ability to systematically investigate the relationship between IDR sequence and molecular function remains limited. For folded domains, the surgical precision with which mutations can disrupt key interfacial residues has enabled the mapping of functionally essential sites at a near-proteome scale^4,5^. In contrast, our ability to design mutations to provide information on disordered proteins has lagged. In some cases, shuffling regions has little effect on function^6–8^, yet even for those same regions, precise mutations that disrupt specific functionally important features can have significant phenotypic consequences.

Pioneering work has paved the way for an alternative path forward: one in which global or local sequence properties can be modulated to investigate sequence-to-function mappings^9–15^. While this approach has been remarkably effective in defining the contours of sequence-ensemble and sequence-function relationships for IDRs, doing so at the scale of tens, hundreds, or thousands of sequences presents a formidable design challenge. At least in part due to this limitation, the explosion in deep learning for protein design that has transformed our ability to engineer function has been largely limited to folded domains^16–19^. Overall, these limitations reflect that fundamentally distinct paradigms for mutation and design are required to tease apart the molecular underpinnings and functional importance of disordered proteins (**Fig. 1A**).

Here, we present GOOSE, a novel computational framework for the rational design of IDRs (**Fig. 1B**). GOOSE enables the rapid design of both *de novo* synthetic disordered proteins and variants of provided sequences, facilitating broad exploration of sequence space (**Fig. S1**). It allows the systematic design of IDRs by sequence properties (amino acid composition, charge, hydrophobicity, charge patterning, etc.), conformational properties (average ensemble dimensions), chemically-specific intermolecular interactions, or indeed, any arbitrary design constraint (*e.g.,* specific 3D conformational ensembles^20^) (**Fig. 1C**). Importantly, for built-in design objectives, GOOSE can generate thousands of sequences per minute with sequence generation speed scaling approximately linearly with sequence length (**Fig. 1D, Fig. S2**). Despite this, GOOSE has no specific hardware requirements, fully democratizing IDR design. In short, GOOSE opens the door to fast and easy library-scale design of disordered proteins to systematically investigate sequence-to-function relationships.

In this work, we illustrate these different design modalities and showcase the power of sequence design to achieve bespoke IDP functionalities. First, we use a FRET reporter to investigate how IDR sequence properties determine ensemble dimensions in living cells. Next, we show how IDR ensemble dimensions relate to subcellular localization and structural sensitivity to cellular perturbations by designing *de novo* disordered proteins with specific ensemble dimensions. Further, we use GOOSE to design disordered proteins that spontaneously self-assemble in cells and enable the targeted recruitment of designed clients, demonstrating our ability to design IDRs with specific intermolecular interactions. Finally, we use GOOSE to design a library of IDR variants to investigate the relationship between sequence and desiccation tolerance in yeast. Taken together, these vignettes highlight the types of investigation GOOSE enables, in the process uncovering insights into how function is encoded in disordered proteins sequence.

## RESULTS

### Investigating sequence-to-ensemble properties in living cells

Substantial work has explored sequence-to-ensemble relationships in IDRs^14^. The same interactions that underlie protein folding also determine transient secondary structure and transient long-range interactions. These conformational biases determine global or local IDR dimensions^21^, accessibility of binding motifs^22^, and change cooperativity between multivalent binding sites^23^.

IDRs can be described in terms of their global ensemble dimensions - the average size occupied by the conformational ensemble (**Fig. 1A**)^24^. Ensemble dimensions can be quantified using various metrics, including the radius of gyration and the end-to-end distance. The ensemble-averaged values of these properties are determined by the balance between attractive and repulsive interactions, which can be both intramolecular (dictated by the protein sequence) and intermolecular (dictated by the surrounding solution environment). Prior *in vitro* and *in silico* work has established core heuristics that inform expected ensemble-average dimensions, yet systematic studies of these principles in live cells are scarce ^14,24–30^.

To interrogate sequence-to-ensemble relationships *in situ*, we measured ensemble-average dimensions for 32 *de novo* IDRs designed using GOOSE in live cells (**Fig. 2A**). All IDRs were sixty residues and designed to enable pairwise sequence comparisons that differed by a single sequence property (**Table S1**). We placed each IDR between two genetically encoded fluorophores forming a Fluorescence Resonance Energy Transfer (FRET) donor/acceptor pair, allowing us to use FRET efficiency (E_f_) as a proxy for end-to-end distance (i.e., ensemble dimensions; **Fig. S3**)^31,32^. If the ensemble is compact, E_f_ is high; if the ensemble is extended, E_f_ is low (see the limitations of the method in *Discussion*). Using these 32 sequences, we assessed how varying sequence properties altered the ensemble-average conformational behavior of IDRs in cells (**Fig. 2A).**

**Figure 2.**
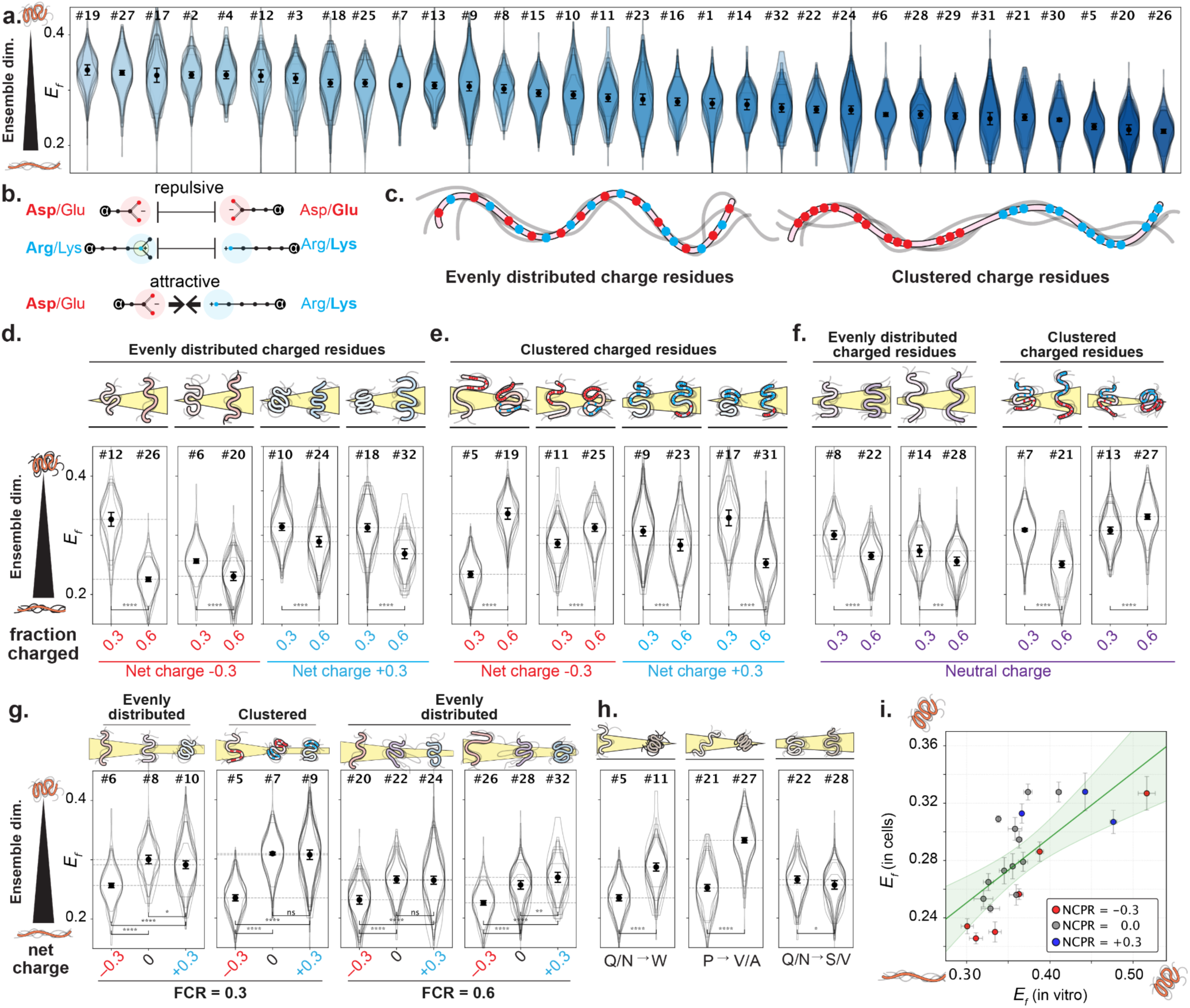
Sequence influences IDR dimensions in live cells. **(A)** FRET efficiencies, E_f_, were obtained from live cell microscopy for all 32 designed sequences (**Fig. S6)** In this and all violin plots in this work, each of the overlaid violins represents a biological repeat with n > 30 cells (**Fig. S3**). The markers and error bars in the center of each violin are the average and standard deviation of the medians of all biological repeats. **(B)** Charged residues of the same polarity are repulsive, while charged residues of opposite polarity are attractive. **(C)** We focus on sequences in which charged residues are either evenly distributed (left) or in which like charges are clustered (right). **(D)** E_f_ for IDRs with evenly distributed charged residues and a net charge of –0.3 or +0.3. **(E)** E_f_ for IDRs with clustered charged residues and a net charge of –0.3 or +0.3. **(F)** E_f_ for IDRs with a net neutral charge, upon increasing the fraction of charged residues from 0.3 to 0.6. **(G)** E_f_ For similarly designed IDR triplets with different charge patterning and fraction of charged residues, systematically titrating the net charge from –0.3 to 0 to +0.3. **(H)** E_f_ for sequences with specific and interpretable non-charged sequence changes, including mutation to aromatics, removal of prolines, and substitution for H-bond donor residues, going from the left to the right panel, respectively. **(I)** Comparison of in cell *vs in vitro* E_f_ values. Residues with a net charge of −0.3 are shown in red and +0.3 in blue. The green line is a linear fit to the data, with the 95% confidence intervals shown by the shaded area. Error bars for in vitro data are obtained from the standard deviation between 3 biological repeats.

Our experiments showed that with few exceptions, most of the rules elucidated *in vitro* hold true under cellular conditions. Focusing initially on sequences with evenly distributed charged residues (**Fig. 2B, C**), we found that enhancing the net charge (either positive or negative) by increasing the fraction of charged residues made ensembles more expanded (**Fig. 2D**). This is in good agreement with *in vitro* work where both the net negative and net positive charges have been found to correlate with chain dimensions^26–28^. Next, we examined how the clustering of charged residues alters ensemble behavior^12^. *In vitro* and *in silico* work has established that charge patterning – the relative clustering of oppositely charged residues – can impact intramolecular interactions and ensemble compaction^12^. As expected, increasing the fraction of charged residues (FCR) for sequences with charge clusters led to ensemble compaction in IDRs with a net negative charge (**Fig. 2E**, left). In contrast, for sequences with charge clusters and a net positive charge, increasing the FCR led to ensemble expansion, an unexpected result that we speculate may be driven by these polycationic IDRs interacting with other cellular components (**Fig. 2E**, right; see *Discussion*). Taken together, our results show that for IDRs with a net charge, both the FCR and the patterning of charged residues can strongly influence IDR dimensions in cells.

We next explored how the FCR of strong polyampholytes (net-neutral sequences) influences their ensemble dimensions. For strong polyampholytes with evenly distributed charged residues, we observed an increase in ensemble dimension with increasing FCR (**Fig. 2F**, left). However, while we expected that increasing FCR would drive compaction in strong polyampholytes with clustered charges, we observed both compaction and expansion (**Fig. 2F**, right). This may be due to the presence of more prolines in expanded sequences (+6 Pro for #7 vs. #21) compared to compact sequences (+1 Pro for #13 vs. #27), suggesting that proline content (driven by chain stiffness and/or the favourable solvation energetics of proline) can also tune intra-chain interactions, consistent with prior work^27,29,33^. Overall, and as with IDRs that possess a net charge, our results show that both the FCR and patterning of charged residues tune IDR dimensions in cells.

Next, we asked how IDR ensembles with equivalent FCR values but different charge signs (negative, neutral, or positive) behaved. We designed sequence triplets with FCRs of 0.3 or 0.6 in which the net charge per residue varied (from –0.3 to 0.0 to +0.3), with sets having either evenly distributed or clustered charged residues (**Fig. 2G**). *In silico* and *in vitro* studies predict that the ensemble dimensions of IDRs with the same charge but opposite signs should be similar^26–28,34,35^. In contrast, our results again showed that net positively charged sequences have similar dimensions to net neutral sequences, and both are far more compact than acidic sequences. This holds true at a FCR of both 0.3 and 0.6 and clustered or evenly distributed charges. Interestingly, the relatively invariant expanded ensembles of negative sequences can be over-ridden by introducing aromatic residues (**Fig. S4**)

To assess the extent to which our conclusions are sequence-intrinsic vs. environment-dependent, we measured 21 of 32 sequences using *in vitro* FRET (**Fig 2I**). This comparison yielded a generally good correlation (Pearson’s r = 0.70 ± 0.04). Pairwise comparisons of the *in vitro* data revealed that, as with the in-cell constructs, increasing the fraction of charged residues with evenly distributed charge patterning expanded ensemble dimension (**Fig. S5A).** Other sequences also displayed expected behavior *in vitro*. For instance, the addition of aromatics led to compaction (**Fig. S5B**). Crucially, sequences that displayed unexpectedly compact ensembles when positive charges were added in-cell showed no significant change or a mild compaction when measured *in vitro* (**Fig. S5C**), supporting the hypothesis that these sequences interact with cellular components^36,37^. Furthermore, sequences with a net negative charge were more compact *in vitro* than in cells (**Fig. 2I**).

Taken together, these experiments identify several differences between *in vitro* and in-cell observations, but our results largely confirm that the same properties that influence IDR dimensions in vitro also hold true in cells. Increasing the fraction of charged residues makes polyelectrolytic IDRs (sequences with a net charge) expand, while this effect is more muted for polyampholytic IDRs (sequences without a net charge). Most notably, negative polyelectrolytes are highly expanded, while polyelectrolytes that contain positive charges are unexpectedly compact.

### Designed ensembles to control IDP behavior in the cell

Having uncovered principles defining sequence-ensemble relationships for our library, we next investigated whether we could detect phenotypic differences in how designed sequences behave in the cell as well as their response to environmental perturbation. We first investigated the relationship between ensemble dimensions and nuclear localization by quantifying the nucleus-to-cytoplasm fluorescence intensity ratio (N/C) (**Fig. 3A**). Previous work has suggested that molecular size influences nuclear import, with smaller proteins diffusing more freely into the nucleus^38,39^. Focusing on constructs that were uniformly dispersed (see *Discussion*), this analysis revealed a general correlation in-line with expected behavior: more compact sequences have a higher nuclear signal, and as ensemble dimensions increase sequences are generally more excluded from the nucleus (Pearson’s r = 0.61 ± 0.09) (**Fig. 3B**). That said, we note that nuclear localization is a complex, muti-faceted phenotype (see *Discussion*).

**Figure 3:**
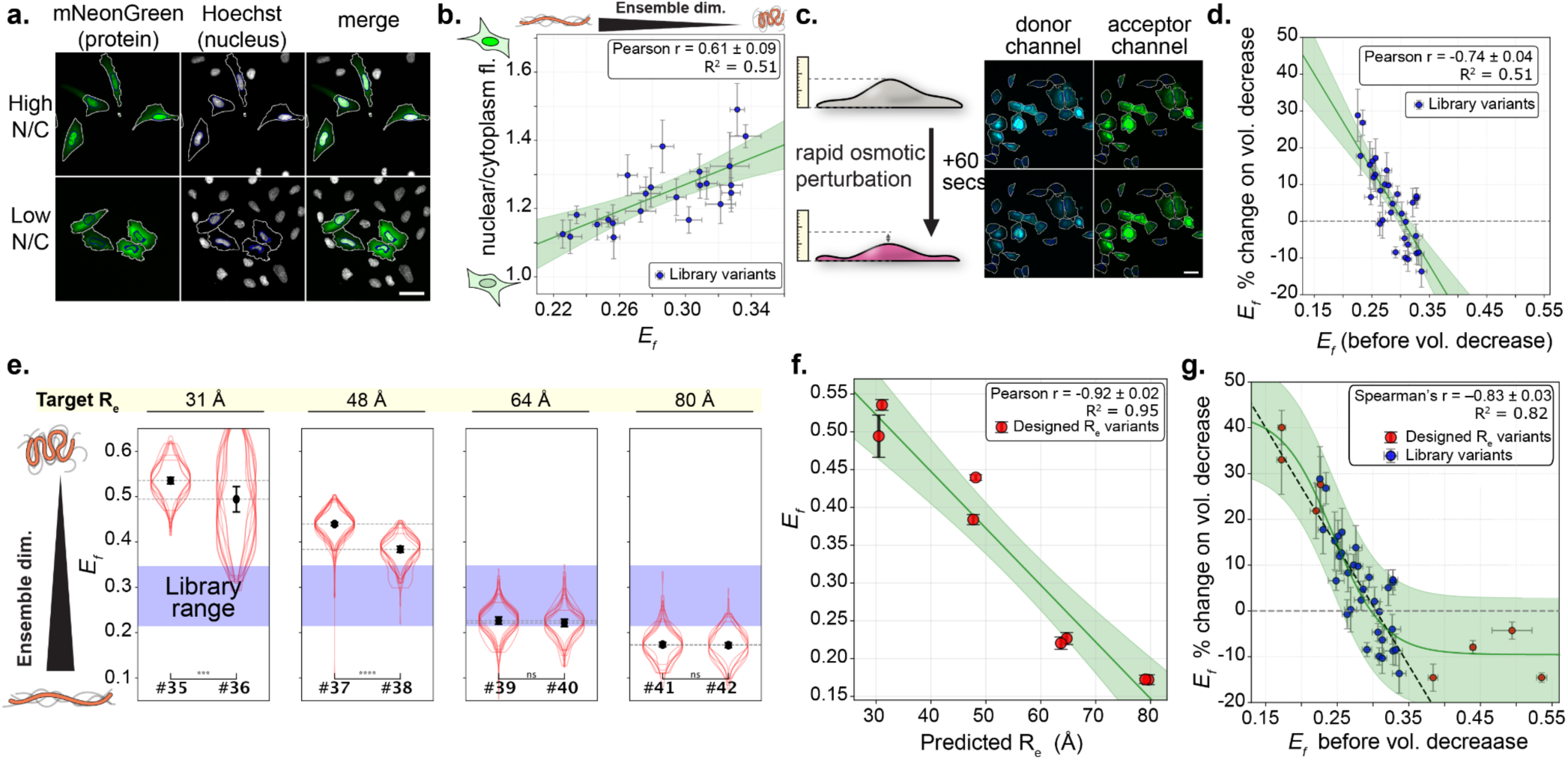
Ensemble dimensions affect nuclear localization as well as conformational response to osmotic shock. (**A**) Representative images displaying differences in nuclear localization between constructs. “High N/C” exhibits nuclear enrichment, while “Low N/C” exhibits nuclear exclusion. Cell mask contours are overlaid on each image, with the cell shown with a white outline and the nucleus with a blue outline. Scale bar = 50 µm. (**B**) Correlation between E_f_ and nuclear/cytoplasm fluorescence intensity ratio per construct. The green line is a linear fit of the data. All shaded green regions in this figure are 95% confidence intervals of the fits. (**C**) Schematic of protocol and representative images of FRET donor and acceptor channels before and after osmotic perturbation with analysis cell mask contours overlaid. All images are acquired 45-120 seconds after perturbation. Scale bar = 50 µm. (**D**) Percent change in E_f_ following hyperosmotic perturbation as a function of E_f_ prior to perturbation. Values above 0 on the y-axis imply ensemble compaction after cell volume decrease, while values below imply ensemble expansion. The green line is a linear fit of the data. (**E**) Violin plots of E_f_ distributions for designed *R_e_* sequences, with design targets above each construct pair. Blue shaded region is the range of median E_f_ values for the original 32 sequence library. Each overlaid violin represents a biological replicate with n > 30 cells, while the markers and error bars are the mean of the medians for the replicates and their standard deviations. Significance is determined between these medians with a student’s two-sided t-test. (**F**) Relationship between E_f_ and *R_e_* as predicted by ALBATROSS. The green line is a linear fit of the data. (**G**) As Fig. 3D, but with designed *R_e_* sequences included. The green line is the data fitted to a variant logistic growth equation, and the dashed black line is the linear fit on library sequences from Fig. 3D.

Next, we sought to investigate ensemble sensitivity to change in cellular volume. We and others have previously shown both *in vitro* and in cells that IDRs exhibit a variety of ensemble-dependent responses to crowding as a function of crowder size^31,32,40^. We therefore wondered to what extent ensemble dimensions dictate crowding sensitivity in a cellular context. To test this we imaged our cells in basal conditions then added mannitol, a membrane impermeable osmolyte, to the imaging media. This causes a rapid cell-volume decrease and concomitant increase in cellular crowding^31,41,42^ (**Fig. 3C**). When comparing cellular FRET efficiencies before and after this perturbation, a strong trend is revealed: expanded sequences become more compact when crowded, while compact sequences show little change (Pearson’s r = −0.74 ± .04) (**Fig. 3D**). The observed compaction of expanded conformations is in line with expectations from macromolecular crowding: as space becomes more restricted, expanded conformations become less favorable, driving ensemble compaction^40,43^. More surprising is the small number of sequences that expand under crowding conditions, possibly due to increased transient intermolecular interactions with binding partners and/or greater concentrations of small-molecule solutes that mask attractive intramolecular interactions^44^. Based on these initial observations, we sought to further investigate the relationship between basal ensemble dimensions and crowding-induced compaction with *de novo* sequences designed to extend our dynamic range of ensemble dimensions.

To directly test whether ensemble dimensions underlie crowding sensitivity, we used GOOSE to design eight additional 60-residue sequences that systematically titrate predicted end-to-end distances from highly compact to highly expanded (target R_e_ (Å) = 31, 48, 64, 80) (**Fig. 3E**, **Table S2**, **Fig S7**). Importantly, these designs substantially expand the dynamic range of ensemble dimensions from those seen in our original library. The ensemble dimensions of these sequences were measured under basal conditions, when we examined the ensemble dimensions of our new sequences, they showed the expected correlation between E_f_ and their design objective dimensions (**Fig. 3F**), establishing our ability to use GOOSE to design IDRs with desired ensemble properties.

Having extended our ensemble dimension space with the novel synthetic IDRs, we tested our eight new sequences using the same rapid cellular crowding protocol. As expected, highly expanded sequences show the greatest compaction upon crowding. In contrast, we observed that when basal ensemble dimensions drop below those of the initial range library, a plateau occurs; beyond this point, increased crowding fails to trigger further sequence expansion (**Fig. 3G**). These results showcase our ability to design sequences that will respond with a large ensemble change when the cell is exposed to a hyperosmotic challenge, as well as our ability to uncover core physical principles of intracellular environmental responsiveness.

Taken together, the correlations we observe showcase the power of designing IDR sequences with specific dimensions. Other phenotypes, including but not limited to diffusion rates, binding affinity, and lifetimes may also be modulated by sequence design and so are a viable target for the systematic sequence design offered by GOOSE. Our results also showcase the continuation of trends when ensemble dimensions are taken to the extreme, into sequence space where chemistries and the resulting ensemble are far outside of the biological norm.

### Rational design of disordered proteins that self-assemble in *S. cerevisiae*

While our designs thus far focused on intramolecular interactions, we next turned to designing for intermolecular interactions. IDRs frequently interact with other biomolecules via sequence-specific (also referred to as site-specific) or chemically specific interactions^1^. Sequence-specific interactions are driven by local motifs in which the precise order of the underlying amino acids is critical, leading to a structured bound state (**Fig. 1A**). Chemically-specific interactions do not depend on a precise sequence but rather on local or global complementary chemistry.

We recently developed a computational approach to predict chemically specific molecular interactions mediated by IDRs (**Fig. 4A**)^45^. This method (FINCHES) allows us to take two disordered sequences and predict a mean-field intermolecular interaction score (ε) between them. When this score is negative, the two sequences are predicted to be attractive for one another, and when the score is positive, the sequences are predicted to be repulsive. We reasoned that we could invert this prediction into a design objective, enabling us to design synthetic sequences with desired interaction properties and opening the door to chemically specific IDR design.

**Figure 4:**
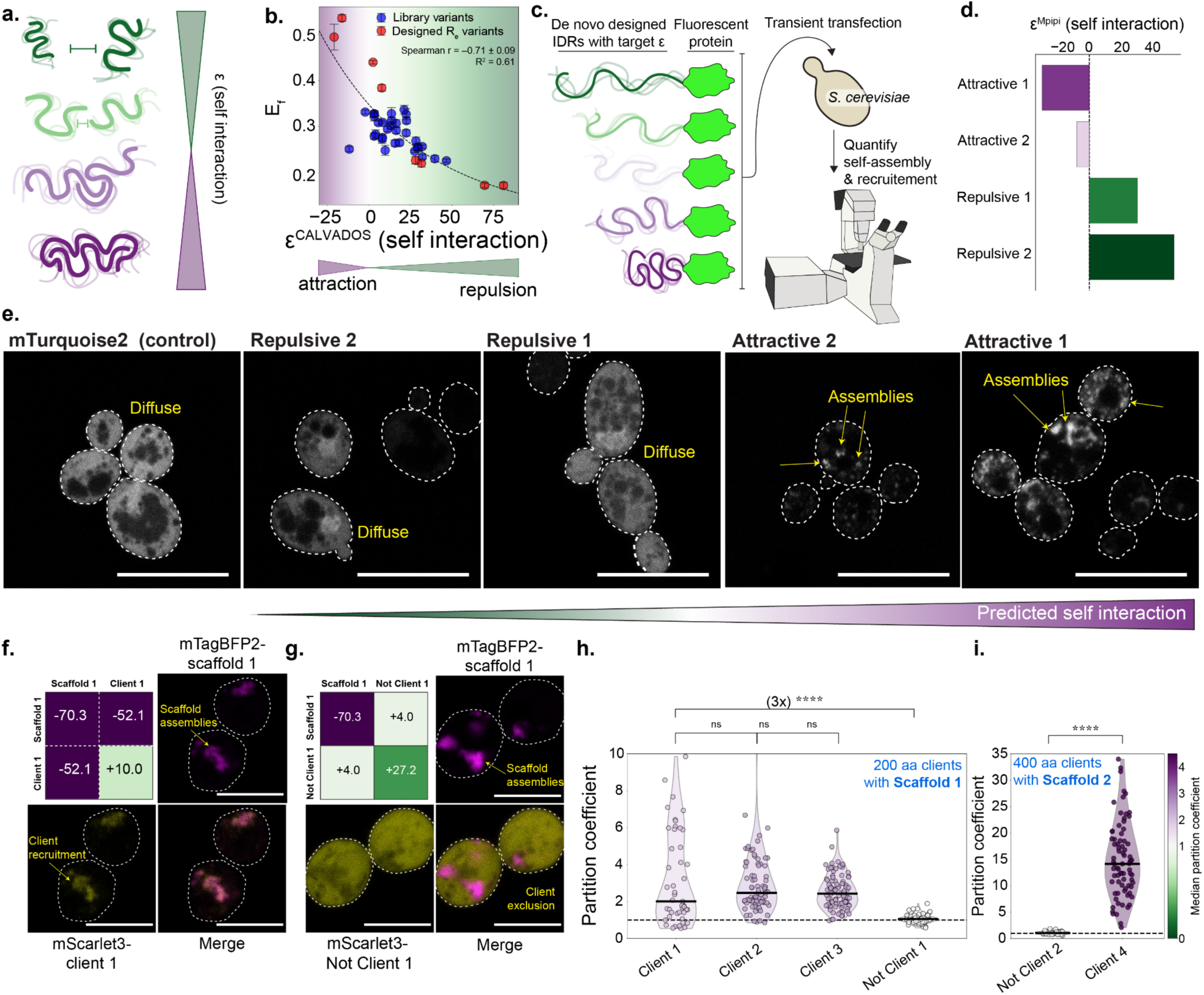
Design of IDPs with specific intermolecular interactions. (**A**) FINCHES enables the calculation of a single, mean-field parameter that captures IDR:IDR-mediated interactions. **(B)** Correlation of homotypic ε value (prediction) and measured E_f_ value (as reported in Fig. 3). More compact chains (higher E_f_) show a greater degree of homotypic attraction (more negative ε). ε was calculated using the CAVALDOS model. **(C)** Overview schematic of experimental setup for quantifying intracellular interaction. **(D)** Self-interaction epsilons for four 400-residue sequences designed to titrate self-interaction from attractive to repulsive. ε was designed for and calculated using the Mpipi model. **(E)** Results of self-assembly designs. The fluorophore (mTurquoise2) is uniformly distributed across the cell, as are the IDRs designed to be self-respulsive (positive ε). In contrast, IDRs designed to be self-attractive (negative ε) form cellular assemblies, with the more self-attractive sequence undergoing greater self-assembly than the less self-attractive one. Scale bar = 10 µm. **(F)** Microscopy images for a 2-component system in which a self-assembling scaffold (Scaffold 1) forms, and a second component was designed to be recruited to that scaffold (“Client 1”). Intermolecular ε calculated using Mpipi. Scale bars = 5 µm. **(G)** Microscopy images for a 2-component system in which a self-assembling scaffold (Scaffold 1) forms, and a second component was designed not to be recruited to that scaffold (“Not Client 1”). Intermolecular ε calculated using Mpipi. **(H)** Quantification of co-localization of 400-residue scaffold (Scaffold 1) and 200-residue clients. Violins show Kernel Density Estimate across partition coefficients and are colored by median partition coefficient. The back line reports the median of each distribution. Individual data points are shown for individual assemblies in different cells. Client 1,2,3 are statistically indistinguishable for one another, whereas Not Client 1 is significantly different from all other constructs (p < 0.001, Pairwise Mann-Whitney U with Holm-Bonferroni correction, see **Fig. S10**). **(I)** Analogous data for a 400-residue scaffold (Scaffold 2) and a 400-residue client. Larger clients show substantially greater partitioning (∼10x), in agreement with expectations from enhancing multivalency. The 400-residue Not Client 2 and 400-residue Client 4 are significantly different from one another (p < 0.001, Pairwise Mann-Whitney U with Holm-Bonferroni correction, see **Fig. S11**).

Biomolecules can engage in homotypic interactions (interactions between multiple copies of the same type of molecule) or heterotypic interactions (interactions between different types of molecules). FINCHES allows us to design sequences with homo- or heterotypic interactions. Focusing initially on homotypic interactions, we used intracellular self-assembly as a proxy for homotypic interactions. We reasoned that IDRs with strong homotypic interactions would be likely to undergo self-assembly in cells, whereas those with repulsive homotypic interactions would remain dispersed. While there are some caveats associated with such an approach (see *Discussion*), this logic has been used to explore naturally-occurring sequences over the last decade ^46–48^.

To establish the validity of using ε as a design criterion in a cellular context, we re-analyzed our designed sequences reported in **Fig. 3** and compared predicted ε values with basal E_f_. Prior work has established a quantitative relationship between IDR dimensions and homotypic interaction strength, driven by an enthalpic equivalence between intra- and intermolecular interactions in long, semiflexible polymers^30,45,49^. Indeed, we find a good correlation between E_f_ and ε (**Fig. 4B**), suggesting that sequence-derived ε values can faithfully report on IDR interactions in a cellular context.

Next, we designed a series of long (400-residue) IDRs with variable homotypic interaction strengths and transformed a fluorophore-tagged version of each into *S. cerevisiae* (**Fig. 4C,D**, **Table S3**). We then examined the subcellular localization of our fluorophore-tagged IDRs and compared them with the fluorophore alone (**Fig. 4E**). The fluorophore-alone control and two self-repulsive IDRs all appeared diffuse in the cell (**Fig. 4E**, left). In contrast, the two sequences designed to be self-attractive showed punctate localization (**Fig. 4E**, right). These results demonstrate our ability to design long, *de novo* IDRs that are expressed, tolerated, and undergo self-assembly in living cells.

While our results in **Fig. 4D** could be interpreted in terms of homotypic interactions, we cannot exclude the possibility that these designed IDRs are co-assembling with other cellular components. Large, disordered proteins designed for chemically specific attractive interactions are likely to be relatively promiscuous^45^. As such, we next sought to test our ability to encode specificity into IDR-mediated interactions by developing a simple two-component system with a basal self-assembling 400-residue “scaffold” IDP, and a series of 200-residue “clients” that would either be recruited to or excluded from the scaffold (**Fig. 4E**). All designed sequences are shown in **Table S4**.

To test our ability to design IDRs with specific molecular interactions, we quantified co-localization of scaffolds and clients in yeast. We first established that our scaffold disordered proteins undergo robust self-assembly in the absence of any clients (as designed) (**Fig. S8**). Next, we co-expressed clients designed to be recruited to (**Fig. 4F**, “Client” designs) or excluded from (**Fig. 4G**, “Not client” designs) scaffold assemblies. These designs followed the expected behavior, confirming our ability to engineer synthetic IDRs with desired chemical specificities. Finally, we reasoned a 400-residue client would show substantially enhanced recruitment compared to a 200-residue client. Designing and testing a 400-residue client, we found ∼10x greater client recruitment, despite the client being entirely diffuse in the absence of the corresponding scaffold (**Fig. S9B)**. Indeed, across all designed clients for scaffold 1 and scaffold 2, we recovered expected partitioning profiles, despite substantial variation in sequence between clients and across the two scaffolds (**Fig. 4H, I, S10, S11)**. Overall, these results suggest that we can employ GOOSE in conjunction with FINCHES to design IDRs with the desired intermolecular interaction profiles, and that these interactions are preserved *in vivo*.

### Library-driven design of disordered proteins that protect yeast from desiccation

Previous sections focussed on the *de novo* design of synthetic IDRs with desired chemical, conformational, or interaction properties. However, a major advantage of GOOSE is its ability to generate variants from template sequences with specific constraints, and to do so at scale. To showcase this capability, we examine the effects of sequence variations on the functional desiccation tolerance conferred to yeast by IDPs. While multiple strategies confer desiccation tolerance in a range of organisms^36,50,51^, the synthesis of protective IDPs is used across all kingdoms of life^52–54^. However, systematic exploration across multiple template sequences and different sequence properties has thus far been limited ^53,55–58^. Here, using several naturally occurring and synthetic desiccation-protective IDPs as templates, we designed a library that systematically varies sequence properties across thousands of variants. By measuring the ability of the resulting sequences to protect yeast from a desiccation, rehydration, and regrowth cycle, this approach enables us to map sequence-to-function for desiccation protection.

To design our library, we chose 11 chemically orthogonal template sequences (**Table S5**). Templates originate from different organisms, including *Hypsibius exemplaris* (tardigrades), *Arabidopsis thaliana* (mouse-eared cress), *Vitis vinifera* (Grape), among four other plant species, as well as several synthetic sequences. For each template, a series of variants was designed that systematically titrates their amino acid composition, amino acid patterning, and global dimensions (**Fig. 5A, Fig. S12**). In total, our library encompasses 2,300 rationally designed disordered protein sequences (**Table S6**).

**Figure 5.**
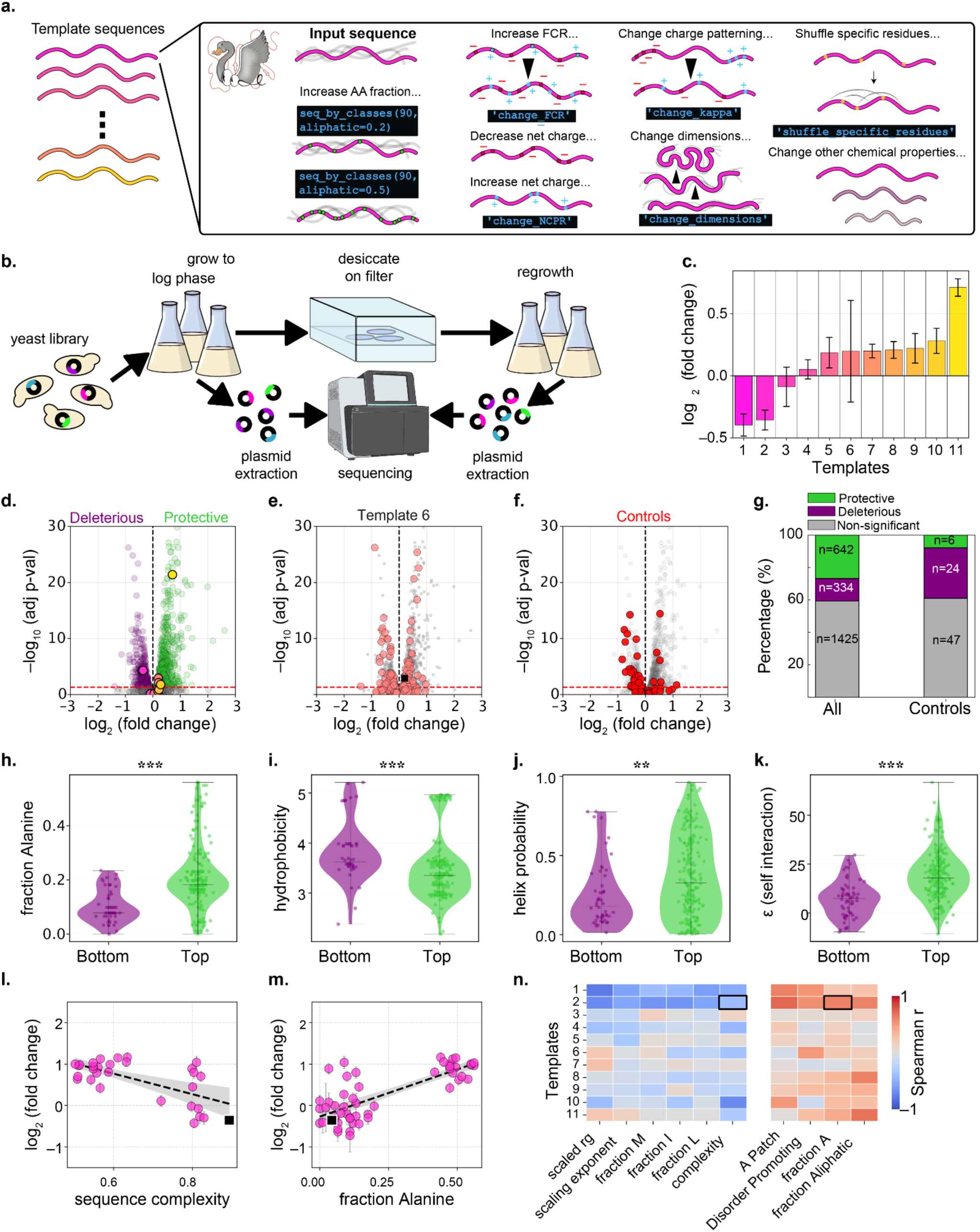
Rational design of desiccation protective IDRs. (**A**) Experimental schematic showing how GOOSE was used as a tool to generate an IDP variant library. **(B)** Schematic representation of the experimental pipeline. **(C)** Template sequences organized by increasing fold-change in reads before and after desiccation and regrowth. Protective sequences have positive log fold-changes. Error bars represent pooled standard error from inverse-variance weighting across three replicates. (**D**) Volcano plot showing the fold change of a given sequence and its false-discovery adjusted p-value. The reported fold changes are averaged across 3 different repeats of the experiment, with each experiment containing 3 biological repeats. The p-value was calculated using a t-test of the weighted average fold changes for each sequence, corrected for multiple testing with Benjamini-Hochberg. Purple and green points are deleterious and protective sequences, respectively. (**E)** Volcano plot as in (D), showing the variations of a single template sequence shown in black. (**F**) Volcano plot as in (D) showing only library internal controls. **(G)** Percentage bar graphs characterizing the extent of protection from all 2300 variants, compared with the internalized controls. (**H-K**) Violin plots characterizing the most protective sequences (‘top’, green) and most deleterious sequences (‘bottom’, purple). A two-tailed Mann-Whitney U test was used to determine whether these groups differed. (**L-M**) Scatterplots of correlations between fold change and specific sequence features: the sequence complexity **(L)** and the fraction of alanines **(M)**. **(N)** A heat map of Spearman’s r for several sequence features and each template. Red colors indicate a positive correlation, blue colors indicate a negative correlation.

To explore how sequence variation influences cell survival following desiccation, the genes were synthesized as a pooled library and cloned into a low-copy-number plasmid with a strong promoter. The plasmid library was transformed into *S. cerevisiae* BY4742 cells to create a pooled cell library that was used for all experiments. The library was grown to mid-log phase, then subjected to desiccation for 8 hours in a humidity-controlled chamber (RH < 10%). Afterwards, cultures were rehydrated and placed in a turbidostat with synthetic complete media lacking Histidine (SC-His), enabling surviving cells to expand while non-viable cells were depleted (**Fig. 5B**). Tracking cell growth through turbidity measurements in the turbidostat revealed a consistently faster growth in library cells compared to an empty vector control (**Fig. S13**).

To reveal the conferred protection of each sequence, plasmids were extracted from the pooled yeast library before and after desiccation and regrowth. Library sequences were amplified from each library and barcoded, allowing us to quantify the abundance of each sequence. The fold change (FC) in normalized read abundance after vs. before desiccation and regrowth allowed us to assess the degree of protection conferred by each sequence: if a sequence was protective, it would increase in abundance (positive FC). If it were deleterious, its abundance would decrease (negative fold change). We found both significantly deleterious and protective sequences in templates (**Fig. 5C**) and across the entire library (**Fig. 5D**).

Focusing on individual templates, sequence variations designed by GOOSE showed that the protective capacity, as measured by the fold change in read abundance, can increase or decrease (**Fig. 5E**, **S14**). As a negative control, the library included sequences with early stop codons, which should display little to no effect in terms of protective capacity (**Fig. 5F**). Of 77 controls, only 6 (8%, compared to 27% for non-control sequences) showed protective effects (**Fig. 5G**), and these were mostly minor.

With the protective capacity of each sequence in hand, we next asked what sequence features promoted (or reduced) protection. Using a combination of sequence analysis and prediction packages (NARDINI+^59^ and sparrow^34^), we calculated or predicted 130 different sequence features for each sequence (**Table S6**). We first note that the majority of sequences with a significant effect were protective and not deleterious (**Fig. 5D, G**). Focusing on the most protective (logFC > 0.5, adjusted p-value < 0.05, n = 216) and most deleterious sequences (logFC < −0.5, adjusted. p-value < 0.05, n = 63), we found features that are significantly different between the two classes across all templates (**Figs. 5H–K, Fig. S15)**. Protective sequences showed significantly higher alanine content (**Fig. 5H**) and lower hydrophobicity (**Fig. 5I**), consistent with reports on functional desiccation-protective proteins^60,61^. Protective variants also showed higher predicted helical propensity (**Fig. 5J)**, a conformational transition well-documented in hydrophilic IDPs during water loss^52,62–65^.

Another additional interesting difference was the higher self-repulsion (positive *ɛ* values) of protective sequences (**Fig. 5K**). This is in contrast to recent observations showing that self assembly of desiccation-protective IDPs can promote desiccation tolerance^66–68^. We note that where self-assembly has been reported, this is driven by specific molecular interfaces that give rise to structured, percolated networks, as opposed to the multivalent interactions that lead to dense, disordered droplets often associated with disordered regions^68^. Our results suggest that an optimum exists - molecular shielding^58^ requires protectants to extensively interact with the cellular milleu, but this interaction may be limited in overly self-attractive sequences. We tentatively suggest a finely tuned balance that maximizes interaction with client molecules and minimizes self-association (which would reduce the effective concentration of available protectants for cellular components) governs this interplay.

To gain a more comprehensive understanding of sequence variations that confer protection, we quantified the correlation between protection and sequence features for each template, excluding sequences within 7% similarity to the template value (**Figs. 5L–M)**. Quantifying the correlation between fold-change and sequence-feature variation for each template (**Fig. S16A**), several patterns emerge: for nearly all templates, increasing disorder, aliphatics, and, specifically, the alanine fraction and patchiness enhanced protective capacity (**Fig. 5M)**, consistent with the overall trends (**Fig. 5H**, **Fig. S15**). Conversely, increasing the fraction of hydrophobic residues broadly correlated with reduced protection (consistent with self-interaction being detrimental, **Fig. 5K**)^58,61^, and decreasing sequence complexity was similarly beneficial ^69^(**Figs. 5M**). However, across nearly all feature variations, we find that the ability to modulate protective capacity is strongly dependent on the sequence context itself. This highlights the disparate ‘designability’ of IDRs: some sequences are ‘fixed’ in terms of ability to protect a yeast cell from desiccation, and no matter what chemistries are changed, there is little to no effect.

Taken together, our results showcase the power of rationally designed IDP libraries to uncover the molecular grammar of IDP function. While our results covered a relatively small library, it can be extended and easily modified to create an iterative pipeline that produces potent functional IDPs tailored to specific functions in targeted environments.

## DISCUSSION

IDRs constitute more than 30% of the human proteome and play critical roles in almost every aspect of eukaryotic biology^1^. Despite this, our understanding of how IDR sequences relate to molecular and cellular functions is only beginning to emerge^14^. A major barrier to investigating how IDR sequences relate to function is the challenge of determining how to iteratively and systematically perturb IDRs in an interpretable manner. Structural biology has been instrumental in our working model of molecular function, in no small part because structural information provides an immediate route to hypothesis-driven investigation. For IDRs, the absence of a ‘reference’ structure from which sequence changes can be mapped to raises a conceptual barrier: how do we begin to start investigating the determinants of IDR function absent a “structural” basis for function?

To address this challenge, we introduce GOOSE, a novel tool for designing *de novo* and variant disordered protein sequences. GOOSE enables the rapid, systematic generation of IDRs that vary built-in design objectives for key sequence features and properties that emerge as important determinants of IDR function. Beyond these built-in methods, GOOSE can also be used alongside arbitrary design objectives to generate IDRs with bespoke sequence properties, as well as to facilitate multi-objective optimization. Across several different systems (*in vitro*, in yeast, in human cells), we illustrate a range of use cases for GOOSE, revealing distinct sequence-encoded functional determinants in the sensitivity of IDPs, their ability to undergo homotypic and heterotypic intermolecular interactions, and their ability to function as protective agents against environmental stress.

Examining sequence-ensemble relationships in cells, we find results consistent with established theory: increasing net charge, the overall fraction of charged residues, and proline residues tend to expand the ensemble dimension (**Fig. 2D,E,H**), whereas adding aromatic residues tends to lead to compaction (**Fig. 2H**). However, for positively charged residues, results are less clear cut. While increasing the net charge of sequences led to expansion in some cases (**Fig. 2D**), in others the addition of positive charge had no effect (**Fig. 2G**) or even led to compaction (**Fig. 2G**). Furthermore, sequences with blocks of positively charged residues became more expanded (**Fg. 2E**) or and more compact (**Fig. 2F**) as the fraction of charged residues increased. Clusters of positively charged residues can act as nuclear localization signals, and many sequences containing them are recruited into nuclear puncta (**Fig. S6**). This is consistent with prior work showing that an abundance of positive charges is sufficient for recruitment into a variety of nuclear subcompartments enriched in RNA^37,70^. Furthermore, our negatively charged sequences were generally more expanded in cells than *in vitro*. These results are largely consistent with a cellular milieu where positively charged biomolecules tend to interact, and negatively charged biomolecules are expanded and more soluble, consistent with a growing body of work across a variety of organisms and systems^36,37,71^.

Our sequences, designed to achieve specific ensemble dimensions, confirm a strong correlation between sensitivity to volume change and ensemble dimensions (**Fig. 3G**). However, we observed a weaker correlation with nuclear localization (**Fig. S17**). This finding is expected: passage through the nuclear pore depends on a combination of factors, including the presence of nuclear localization signals, chemical miscibility with the nuclear pore environment^72^, ensemble dimensions, and the availability of soluble protein for nuclear uptake^73^. While one of the constructs, designed to be compact (#35), shows substantial nuclear enrichment, another (#36) forms cytoplasmic assemblies, likely limiting the pool of soluble protein available for nuclear import (**Fig. S6)**. These results highlight the complex interplay among competing factors, which can obfuscate mechanistic insights when physical models for the underlying processes are lacking.

It is important to note that for all FRET experiments, the presence of terminal fluorescent proteins (FPs) certainly plays a role in the final conformations of the ensemble - whether this is through IDR:FP interactions or through self-association. Nonetheless, we believe that these interactions do not dominate our conclusions: all designs were based solely on the IDP sequence. Overall agreement with predicted IDP dimensions (**Fig. 3E**), a high correlation with self-ε (**Fig. 4b**), and E_f_ shifts seen in pairwise sequence comparisons that follow expected behavior (**Fig. 2**) indicate that these interactions are not a major determinant of the observed signal.

We examined our ability to design synthetic IDPs that undergo self-assembly *in vivo*, and recruit client molecules to those assemblies. These assemblies are prime examples of biomolecular condensates – non-stoichiometric cellular bodies that concentrate specific biomolecules and exclude others; however, we emphasize we make no claims regarding either their composition or material state^74^. Although self-assembly is a convenient readout of homotypic or heterotypic protein-protein interactions, it is not without limitations. We acknowledge that our assays examining self-assembly were conducted in the intracellular environment, where many other potential interacting partners exist beyond our designed IDRs. Although we demonstrated five designs that successfully formed self-assemblies (including Scaffold 1 and Scaffold 2), it is important to note that designing an IDR to be self-attractive does not guarantee assembly formation *in vivo*. Furthermore, off-target interactions with other cellular biomolecules cannot be ruled out. Similarly, our colocalization experiments are not without limitations. As one might expect, these designs proved to be more challenging than simple self-assembly. Specifically, because the clients had variable self-interaction strengths, two of the four designed clients were capable of self-assembly even in the absence of the scaffold (**Fig. S9)**. Although these designs still showed significant co-localization with their respective target scaffold, we cannot rule out that both scaffold and clients are co-localizing with an existing cellular condensate (even if this were the case, the clients would still co-localize as intended). However, two of our four client designs were diffuse in the absence of the scaffold protein, which gives us confidence that these designs showed only punctate localization patterns due to recruitment to their target scaffold.

Beyond self-assembly, we also leveraged GOOSE to explore sequence designability for desiccation protection. The ability to alter sequence protection varied markedly across templates (**Fig. 5N, Fig. S16**). Templates of synthetic origin or unrelated to desiccation protection (templates 1 and 7) showed little designability - protection was low and scarcely improved regardless of sequence variations introduced. In contrast, desiccation-related templates (templates 2, 4, 6, 8–11), known to confer protection from desiccation in other organisms (**Table S5**), showed greater protective capacity and more tunable sequence-function relationships. Notably, these already protective templates could be further improved by sequence variation (**Fig. S14**), suggesting that even naturally-protective sequences have not yet reached a compositional optimum for protection or can be further optimized in the exogenous yeast environment. Importantly, there were very few sequence modifications that universally enhanced protection across all templates; the optimal path to increased protection was template-dependent. The trends that did show cross-template uniformity (increasing aliphatics and specifically alanines, decreasing hydrophobicity, and reducing sequence complexity) align closely with the defining compositional properties of desiccation-protective IDPs^61^.

It is notable that the overall effect measured as fold-change in read counts before and after desiccation and regrowth may appear modest - a factor of 8 at most, with many sequences showing fractional values. This reflects our experimental design, in which regrowth is intentionally limited to ∼15–20 hours following desiccation to prevent sequence drop-out from the library (**Fig. S13)**; prolonging regrowth would drive fold-changes upward. Accordingly, the more meaningful metric is statistical significance, supported here by 11 biological replicates collected across 3 independent experiments.

Knowing which sequence has this enhanced designability seems, at least for desiccation protection, difficult to predict a priori, but it would help future research and the generation of new libraries for improved desiccation tolerance. Using GOOSE and a pooled experimental design to generate variants successfully enables us to pin down this designability and offer explicit strategies for further improvement. Furthermore, this pipeline can be used iteratively to improve a given phenotype. This is a powerful approach that can both uncover sequence-to-function relationships and assist in the design of *de novo* IDPs with strong functional effects.

Work from several groups has provided exciting new tools focussed on the design of disordered proteins^75,76^. These methods include using iterative simulations^77,78^ or machine-learning-based approaches focussed on ensembles, interactions, or generative design via language models^79–85^. We see these methods as complementary (or even synergistic with) to GOOSE; while these approaches enable the generation of synthetic “natural-looking” IDRs, they are not designed for the generation of sequences that systematically titrate across human-interpretable features for hypothesis testing. Further, given GOOSE’s SequenceOptimizer, it seems emerging deep learning methods can be combined with GOOSE to turn predictive methods into design objectives, as reported recently^20^.

For folded proteins, protein design has largely focussed on generating proteins with a desired 3D structure and, more recently, on desired molecular interactions with specific partners. Design pipelines often involve a hierarchical approach in which hundreds of thousands of initial sequences are generated and screened *in silico*, thousands progress to more rigorous computational validation, and tens are expressed and subjected to downstream experimental characterization. This pyramiding reflects the fact that designing stable, soluble folded domains requires cooperative interactions among many residues and regions. For a folded domain to function – regardless of what it does – correct folding and solubility are unavoidable prerequisites. Given that functionally equivalent folded domains from across the kingdom of life often (though not always) are sequence-similar, we (and others) subscribe to a model whereby “structure” encodes a tight coupling between sequence and function, with even small perturbations in sequence that disrupt structure having devastating consequences for function.

For IDRs, a different paradigm emerges. There is no unifying feature that underlies IDR function. Similarly, when assessed using alignment-based metrics, IDRs often appear poorly conserved, reflecting the empirical observation that many sequences can appear equally “functional” across different systems^6,86,87^. Taken at face value, this suggests that, for a given functional outcome, the design space for IDRs is large. As such, if sequence-to-function mapping for folded domains involves a hypothetical fitness landscape with one (or a small number) of deep wells, analogous landscapes for IDRs involve broad, roaming hills and valleys. In this analogy, the small, incremental sequence perturbations that are so informative for exploring folded-domain function may be poorly suited to exploring this landscape. In contrast, we propose that investigating sequence-to-function relationships by systematically perturbing many sequences is more fruitful, at least initially. Rather than testing a few top candidates, we propose that IDR sequence-to-function relationships require large, high-throughput experiments that map these landscapes across tens, hundreds, or thousands of sequences.

One route to do this may be to use evolutionary-scale large language models that have implicitly learned coevolutionary relationships between disordered regions, enabling large sets of “synthetic paralogs” to be generated and tested^82,88^. A non-overallping and complementary strategy is presented in GOOSE, in which users define their own hypotheses about which sequence features matter and then design large libraries capable of simultaneously testing many competing hypotheses in parallel. By enabling the design of large (10,000+) sequence libraries in minutes to hours, GOOSE provides both the speed and the capabilities to match high-throughout genotype-phenotype mapping. Looking forward, this opens the door to broad-reaching investigations of IDR function in natural proteins, as well as a wide range of biotechnological applications.

## METHODS

### About GOOSE

GOOSE (Generate disOrdered prOtiens Specifying propErties) is a software package for the design of disordered protein sequences. The sequences used in this manuscript were designed using the Python (version 3.9+) package GOOSE (https://github.com/idptools/goose). GOOSE uses sparrow (https://github.com/idptools/sparrow/) to calculate sequence properties. Ensemble predictions used for the design of IDRs with a desired radius of gyration or end-to-end distance use ALBATROSS, as implemented in sparrow^34^. ALBATROSS is a deep-learning tool for predicting ensemble-average IDR dimensions directly from sequence and was parameterized based on coarse-grained simulations performed with a modified variant of the Mpipi model^89^. GOOSE uses FINCHES to calculate predicted intra or intermolecular interactions, where FINCHES makes use of both the Mpipi and CALVADOS forcefields^45,89,90^.

GOOSE has two general design approaches: pre-defined generation and variation functionality and sequence optimization. Both approaches allow the creation of fully synthetic sequences or starting from a sequence of interest (**Fig. S12**). For pre-defined functions, fully synthetic sequence generation allows you to specify various properties (hydrophobicity, FCR, NCPR, kappa), fractions of amino acids, average ensemble properties, and patterning properties (ex. how symmetrically positioned aromatic amino acids are). For pre-defined functions for variant generation (when you start with a sequence of interest), GOOSE has eighteen different types of variants that can broadly be categorized as: 1) shuffling variants - different approaches to shuffle a sequence or subsequence, 2) asymmetry variants - change how symmetrically an amino acid or class of amino acids are positions in a sequence, 3) property variants - change individual properties (ex. FCR, hydrophobicity) while keeping others constant while also allowing keeping different aspects of sequence chemistry constant. The sequence-optimization approach extends GOOSE’s original functionality. Sequence optimization uses the SequenceOptimizer class. This has predefined functionality, but is extensible: any Python function that returns a numerical value can be used with the SequenceOptimizer class. FINCHES-based sequence design is built in via the SequenceOptimizer. Although the sequence-optimization approach is more flexible than predefined GOOSE functions, it is slower to generate sequences (typically still on the order of seconds per sequence).

Because of its speed and flexibility, GOOSE is poised to facilitate the rational design of anything from small numbers of sequences to libraries of thousands for the systematic investigation of sequence-ensemble and sequence-function properties. A key feature of GOOSE is its use of the metapredict (V3) backend to enable rapid, accurate assessment of disorder propensity in designed sequences. Developing a fast and accurate disorder predictor (1000s seconds/sequence with state-of-the-art accuracy^91^) was essential to enable high-throughput library design.

GOOSE is open source and can be used as a Python library or within a Google Colab notebook (https://colab.research.google.com/drive/1U9B-TfoNEZbbjhPUG5lrMPS0JL0nDB3o?usp=sharing). We provided extensive documentation (https://goose.readthedocs.io/en/latest/index.html), which is not reproduced in this supplementary information due to length but can be readily accessed through the web.

Functionally, GOOSE relies on a stochastic design algorithm that enables it to generate unique sequences even when numerous sequence properties are specified. This is true for both the predefined functions and for the SequenceOptimizer class. Generally, fully synthetic sequence generation starts with the creation of a ‘base sequence’ that comes close to satisfying user-specified input parameters. From here, various functions are used to fine-tune the sequence so that its parameters match the input parameters. Then, optimization functions are used to minimize sequence disorder while satisfying any sequence parameter constraints. Finally, the sequence is checked for predicted disorder using Metapredict V3. GOOSE is in active development, and new features will be added regularly. The version associated with this manuscript is version 0.2.5 at the time of submission.

### Sequence Property Library Design Rationale

The sequences in this paper that examined the impact of sequence properties on ensemble dimensions were generated by specifying “sequence properties” within specific ranges, enabling maximal pairwise comparisons in which only one property was varied between pairs of sequences. In particular, sequences were designed with the following quantized sequence properties: NCPR of –0.6, –0.3, 0.0, +0.3, +0.6, FRC of 0.0, 0.3, or 0.6, Kyte-Doolittle hydrophobicity of 1.0 or 3.0 (on a 0-to-9 scale), and kappa [κ] (a measure of charge patterning) was set to be between 0.05 and 0.22 (low-to-average, depending on sequence composition) and then above 0.5 for highly clustered sequences. The quantization of hydrophobicity (and 1.0 or 3.0) was selected for two reasons. Firstly, keeping hydrophobicity low reduces the risk that our synthetic IDRs trigger the unfolded protein response. Secondly, because hydrophobicity is intrinsically coupled with FCR, enabling independent variation of FCR and hydrophobicity, it required lower hydrophobicity scores to accommodate highly charged sequences. Given the scope of sequence space for 60-residue disordered proteins and the relatively low-throughput experimental characterization employed here to ensure high-quality data are reported, we opted to approach our design problem by designing sets of pairs of sequences. Each pair enables the specific comparison of one sequence parameter by holding the others fixed while varying a single parameter (e.g., net charge, hydrophobicity, etc.). By designing our library to multiplex distinct hypotheses, the same sequences could be members of multiple pairs, enabling us to systematically test a collection of hypotheses with a relatively low number of sequences.

### Sequences Designed by Dimensions

To further explore the relationship between basal ensemble dimensions and IDR sensitivity to changes in cellular dimensions, we leveraged GOOSE to generate sequences across a wide range of predicted dimensions. These designs used a simple pre-defined function in GOOSE called “seq_by_re”. For these designs, all that we specified was the sequence length and then the objective end-to-end distance in angstroms. Apart from that, the designs were not constrained by any specific sequence properties.

### LEA Sequence Library Design

Because we did not know which sequence properties would influence the function of each input LEA protein sequence *a priori*, we used a simple strategy for the LEA library design. Specifically, we used the many variant functions in GOOSE and designed variants of each starting sequence to span a wide range of sequence properties, with some properties altered and others held constant (**Fig. S12**). Each variant was based on a starting LEA sequence. The specific types of variants generated were: 1) “constant_properties” variants where FCR, NCPR, hydropathy, and kappa are held constant in the generated variants, 2) “constant_properties_and_class” the same as (1) except the generated sequence also has the same number of each amino acid by class as the starting sequence, 3) “constant_properties_and_class_by_order” the same as (2) except the order of the amino acids by class is also retained in the generated variant, 4) “constant_residues_and_properties” variants where the FCR, NCPR, hydropathy, and kappa are all held constant and a specified subset of residues (by class) is also held constant in position and number, 5) “shuffle_specific_regions” variants where specific subregions of the sequence are shuffled while others are left constant, 6) “change_residue_asymmetry” variants where different classes of amino acids were redistributed to be more or less asymmetrical across the sequence, “change_hydropathy_constant_class” variants where the average hydrophobicity is altered but the amino acids by class and position are kept constant, 7) “change_fcr_minimize_class_changes” variants where the FCR is increased or decreased and changes to amino acids by class are minimized, 8) “change_ncpr_constant_class” variants where NCPR is increased or decreased and everything else is held constant, 9) “change_kappa” variants where the asymmetry of the distribution of oppositely charged residues is increased or decreased and everything else is held constant, 10) “change_dimensions” variants where the end-to-end distance was titrated while changes to the sequence were minimized, 11) FINCHES variants where the self-to-self epsilon value as calculated by FINCHES was increased or decreased while minimizing changes to the sequence. For each type of variant, between 6 and 12 sequences were generated for each input sequence. The specific number of variants for each variant type per input sequence varied slightly because some variants require specific residues to be generated. For example, we did not generate variants that titrated the distribution of prolines across the sequence when the input sequence did not contain any prolines. After all variants were generated, they were checked for redundancy to ensure that no two sequences were the same.

### Fluorescence reporter constructs (FRET imaging)

Our fluorescence reporter construct places the disordered protein sequences from our library (**Table S1, Table S2**) between an N-terminal mTurquoise2 FRET donor and a C-terminal mNeonGreen acceptor ^31^. Genes for each IDR were obtained from GenScript and ligated between the two fluorescent proteins using 5’ SacI and 3’ HindIII restriction sites in a pcDNA3.1(+) backbone.

### Mammalian Cell culture

U-2 OS cells were cultured in Corning-treated flasks with Dulbecco’s modified Eagle medium (Gibco Advanced DMEM:F12 1X) supplemented with a final concentration of 10% FBS (Gibco) and 2.5 mM L-glutamine (Gibco). For live-cell microscopy experiments, 8,000 cells were plated in each well of a µ-Plate 96 Well Black treated imaging plate (Ibidi) and allowed to adhere overnight (∼16 hours) before transfection. Cells were incubated at 37 °C and 5% CO_2_. Before transfection, the media was replaced with new, warmed media. XtremeGene HP (Sigma) was used to transfect FRET construct plasmids into U-2 OS cells as per the manufacturer’s protocol. Cells were then incubated at 37 °C and 5% CO_2_ for 24 hours.

### Live-cell Microscopy (FRET imaging)

Images were taken on an Olympus IX83 microscope equipped with a laser-guided autofocus system and using a 20X 0.8 NA dry objective. Excitation light was produced with a Spectra X light engine (Lumencor, Beaverton, OR), and data were collected on two linked ORCA-fusion Gen-III sCMOS cameras (Hamamatsu Photonics, Shizuoka, Japan). For donor, acceptor, and direct acceptor channels, cells were imaged at room temperature before and after perturbation with 5 ms exposure times and 2.5% LED intensity. Imaging was performed by exciting mTurquoise2 at 438 nm (438/29) for donor and acceptor channels, mNeonGreen at 511 nm (511/16) for the direct acceptor channel, and Hoechst dye at 390 nm (390/22) for the nucleus. The emitted light was passed downstream through a triple-bandpass dichroic (467/24, 555/25, 687/145), then split and directed to the two linked cameras using a 510 nm downstream dichroic housed in a Gemini Optical Splitter (Hamamatsu Photonics).

Cells were stained with Hoechst 33342 (Thermo Scientific) to visualize nuclei by adding Hoechst stock (20 mM) to PBS (Gibco) at a ratio of 0.25 µL to 1.5 mL. Shortly before imaging, 250 µL of this solution was added to each well and incubated for 5 minutes, then washed with fluorobrite, after which a final volume of 250 µL Fluorobrite (Gibco) was added. The imaging process was started ∼15 minutes after the final addition of Fluorobrite.

### Osmotic perturbation imaging (FRET imaging)

Osmotic perturbation solution of 1.8 M mannitol was prepared by dissolving 9.84 g of mannitol (Acros Organics) in a final volume of 30 mL Fluorobrite (Gibco) shortly before imaging. Following imaging in isoosmotic fluorobrite media, 50 µL of the mannitol solution was added to four adjacent wells using a multichannel pipette to achieve a +250 mOsm challenge (550 mOsm total). To ensure mixing, the solution was pipetted up and down 10 times, after which the four wells were imaged. This process was then repeated for all wells in the plate. Imaging for each group was completed between 45 seconds and 2 minutes after perturbation, depending on the well in the imaging order.

### Image Analysis (FRET imaging)

Images were analyzed using a custom Python script (**Fig. S18**) available on the associated GitHub repo. Cell masks were created by segmenting using CellPose^92^ with default settings on the FRET donor channel. A nucleus mask was created by thresholding using Otsu’s method on the Hoechst channel. Image sets before and after perturbation were registered to each other (without the Hoechst channel) using a rigid registration algorithm^93^. A constant value for each channel, determined from the mean values of empty wells, was then subtracted (except for the Hoechst channel) from all pixel values to remove background effects at low fluorescence intensities from the ratiometric FRET efficiency metrics. Areas where background subtraction caused signal values to approach 0 are removed from masks and not counted in calculated metrics. Hoechst masks are labelled to their corresponding overall cell mask, and small holes in the Hoechst mask are filled in.

From these finalized masks, a variety of metrics are calculated for each cell. These include mean intensities for the FRET donor, FRET acceptor, and direct acceptor, for the whole cell, the nucleus, and the cytoplasm. Cell FRET efficiencies before and after perturbation (E_f,basal_ and E_f,perturbed_, respectively) were calculated by E_f_ = *F_A_* / (*F_A_* + *F_D_*). Here, *F_D_* is the mean intensity of the donor, and F_A_ is the mean intensity of the acceptor following fluorescence bleedthrough and cross-excitation corrections (**Fig. S19**). E_f,perturbed_ channels were additionally corrected to account for total fluorescence intensity changes observed after perturbation (**Fig. S20**).

The FRET efficiency change due to osmotic perturbation was calculated as ΔE_f_ = E_f,perturbed_ - E_f,basal_. Data is then filtered to remove dead cells, strange morphologies, missing in after images, etc. All filters used are displayed per construct (**Fig. S21**), and include: cell area, nucleus area, cell circularity, ratio between nucleus and cell areas, direct acceptor maximum, and direct acceptor intensity ratio after and before perturbation. Cells are then further filtered on the mean intensity of their direct acceptor channel to avoid intermolecular FRET effects (**Fig. S22**).

### Statistical Analysis (FRET imaging)

The statistical analysis of all experimental data was performed using the SciPy library in Python. Experiments were performed in 96-well plates across multiple cell passages and days, with each well plated and transfected individually. We therefore treated each well as a biological replicate of the experiment. Therefore, the median values for all cells measured per well were used to generate a single violin plot. We excluded wells that contained fewer than 30 cells. The standard deviation and average values were calculated from the medians of all wells from each experimental condition. To assess significance of the differences between two constructs, a two-sided Student’s t-test was performed between all medians of the two constructs.

### Bioinformatic analysis of FRET construct sequences

Bioinformatic sequence analysis for metrics such as NCPR, FCR, kappa, etc. was performed using localCIDER and sparrow (https://github.com/idptools/sparrow/). Disorder prediction was performed using metapredict (V2-FF).

### Bacterial expression and purification of FRET constructs

For *in vitro* expression and purification, FRET constructs were cloned into a pET-28a(+)-TEV backbone. BL21(DE3) *E.coli* cells (New England Biolabs) were transformed with these plasmids according to the manufacturer’s protocol and grown in lysogeny broth (LB) medium with 50 μg/ml kanamycin. Cultures were incubated at 37 °C while shaking at 250 rpm until an optical density 600 of 0.65 was reached (approximately 5 hours), then induced with 1 mM isopropyl β-d-1-thiogalactopyranoside and incubated for 18 hours at 20 °C while shaking at 250 rpm. Cells were collected by centrifugation for 30 min at 3,000 x g, and the supernatant was discarded. The remaining pellet was lysed in lysis buffer (50 mM NaH_2_PO_4_, pH 8.0, 0.5 M NaCl) using a QSonica Q700 Sonicator (QSonica). The lysate was centrifuged for 30 mins at 21,100 x g, the supernatant was collected, and Ni-NTA agarose beads (Thermofisher) were added to the lysate. After incubating with lysate at 4 °C for 15 mins, the beads were centrifuged at 18,000 x g for 15 mins, and the flow through discarded. Ni-NTA beads were washed two times with a wash buffer (50 mM NaH_2_PO_4_, pH 8.0, 0.5 M NaCl, 20 mM Imidazole). The FRET construct was eluted with 50 mM NaH_2_PO_4_, pH 8, 0.5 M NaCl, and 250 mM imidazole, and buffer exchanged into protein buffer (20 mM sodium phosphate buffer, pH 7.4, 100 mM NaCl). Constructs were further purified using Amicon Ultra-0.5 mL Centrifugal Filters (MilliporeSigma), with a 50 kDa cutoff filter. The purified FRET constructs were divided into 100-μL aliquots, flash-frozen in liquid nitrogen, and stored at −80 °C in protein buffer. Protein concentration was measured after thawing and before use using UV–Vis absorbance at 506 nm (the peak absorbance wavelength of mNeonGreen). Protein purity was assessed by sodium dodecyl-sulfate polyacrylamide gel electrophoresis (SDS-PAGE) after thawing and before use. To verify FRET construct sample integrity and fluorophore stoichiometry, we measured the absorbance spectrum of each sample tested before each FRET assay. We report only samples that displayed an absorbance ratio 506/434 (peak mNeonGreen absorbance / peak mTurquoise2 absorbance) of 2.5 ± 0.5, a reasonable ratio given the difference in the molar extinction coefficients of mTurquoise2 and mNeonGreen (34,000 L mol^−1^ cm^−1^ and 116,000 L mol^−1^ cm^−1^).

### *In vitro* FRET experiments

In vitro FRET experiments were conducted in black, flat-bottom 96-well plates (Thermo Fisher) using a CLARIOstar plate reader (BMG LABTECH). Protein buffer and purified protein solution were mixed in each well to a final volume of 150 μl, containing the buffer and the FRET construct at 1 μM protein. Fluorescence measurements were taken from the top of the plate, at a focal height of 6.8 mm, with gain fixed at 1,200 for all samples. For each FRET construct, three biological replicates were obtained from different expressions, with 3 technical replicates per expression performed. Fluorescence spectra were obtained for each FRET construct by exciting the sample in a 16-nm band centered at λ = 420 nm, with a dichroic at λ = 436.5 nm, and measuring fluorescence emission from λ = 450 to 600 nm, averaging over a 10-nm window moved at intervals of 0.5 nm. Base donor and acceptor spectra were obtained using the same excitation and emission parameters on solutions containing 1 μM mTurquoise2 or mNeonGreen alone.

### Calculation of *in vitro* FRET efficiencies

The FRET efficiency (E_f_) of each FRET construct in each solution condition was calculated as described previously^31^. Briefly, this was done by linear regression of the fluorescence spectrum of the FRET construct with the spectra of the separate donor and acceptor emission spectra in the same solution conditions (to correct for solute-dependent effects on fluorophore emission). E_f_ was calculated using the following equation.

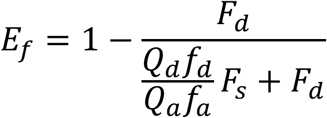

where *F_d_* is the decoupled donor fluorescence, *F_s_* is the decoupled acceptor fluorescence, *f_d_* is the area-normalized donor spectrum, *f_a_* is the area-normalized acceptor spectrum, *Q_d_* = 0.93 is the quantum yield of mTurquoise2 and *Q_a_* = 0.8 is the quantum yield of mNeonGreen ^94^.

### Yeast desiccation library

The sequence designs were reverse-translated into DNA sequences. To ensure proper sequence coverage and distribution across our sequence library, we developed a simple Python package that allows for simultaneous optimization of codon usage and GC content. With this, we were able to generate DNA sequences with optimized *S. cerevisiae* codon usage with an average GC content of ∼40% and a standard deviation of GC content of 0.76% across the library. This tool is publicly available on GitHub at https://github.com/ryanemenecker/kea. The resulting DNA sequences were synthesized on chip (Twist Bioscience) and cloned, using 15-bp homology regions at the 5’ and 3’ ends, into a low-copy-number plasmid with a strong promoter via in-cell homologous recombination^95^. The DNA library was obtained from Twist Biosciences. Each DNA sequence included 12 base pairs of homology to the plasmid backbone at both the 5’ and 3’ ends (5’- ACTAGTGGATCC … CTCGAGTCATGT -3’), and amplified in parallel 16 times in 25 μl reactions with NEB Q5 polymerase in 10 cycles to extend the homology to the backbone (forward primer: 5’- TCTAGAACTAGTGGATC -3’; reverse primer: 5’-ACTAATTACATGACTCGAG -3’). The library was cleaned up using a 1.8x ratio of Magmax beads to PCR product, and the clean DNA was quantified with a Qubit 2.0 dsDNA BR assay, and specific amplification was confirmed on a 3% Agarose gel. The backbone plasmid p413GPD-hTPI was purchased from Addgene and was linearized for cloning by amplification with NEB Q5, after a preliminary digestion with BamHI-HF (NEB). It was amplified in 24 cycles (forward primer: 5’- CACTAGTTCTAGAATCCGTCG -3’; reverse primer: 5’-TCGAGTCATGTAATTAGTTATGTC -3’). The amplified product was cleaned with a Monarch NEB PCR and DNA Cleanup kit, then digested with FastDigest DpnI from Thermo Fisher Scientific in Green Buffer for direct loading onto a 1% Agarose gel. The amplified product was excised from the gel with a Monarch Gel Extraction kit.

To ensure single or low plasmid copies in yeast cells and high expression levels, the p413GPD backbone contained a CEN6/ARS4 integration site and a GPD promoter. Transformation was carried out in yeast by electroporation^96^ and the copies of dsDNA of library:backbone were in a 5:1 ratio to ensure an abundance of insert. Lower library:backbone ratios resulted in fewer transformants. Eighteen transformations were conducted in parallel to guarantee there were enough transformants for the library size, 10x coverage. To achieve this, we plated 200 histidine-deficient agar plates, each with approximately 400 colonies, and scraped a total of ∼80,000 colonies into a 20% glycerol stock.

The plasmid library was transformed into BY4742 yeast cells, and the resulting library was used for experiments, with at least 6 × 10^6^ cells to ensure proper coverage. For each experiment, the cells were grown in histidine-deficient synthetic complete (SC-His) media to select for library expression. We ensured that cells did not experience any stress prior to desiccation by maintaining the culture in log phase using turbidostats. This prevents the yeast from biochemical priming, which confers natural desiccation protection^97^ but would obfuscate the protective effect of the IDPs. The yeast were then subjected to desiccation by pipetting them on nitrocellulose filter paper for 8 hours in a humidity-controlled chamber with low relative humidity (RH < 10%). Following desiccation, each filter paper was rehydrated in SC-His under shaking, and the yeast culture extracted from the filter paper was allowed to grow in a turbidostat, ensuring the culture remains in logarithmic growth for 24 hours (**Fig. S13)**. This regrowth guarantees that cells that did not survive desiccation are depleted, while surviving cells proliferate. The experiment was run at least as triplicate for every experimental repeat, and samples were taken from each replicate before and after DR. Plasmids were extracted from each sample, the variable region of the sequence amplified, and the amplicons were prepared for sequencing. As a control, we compared the pooled library culture to yeast containing an empty (non-expressing) vector under the same promoter and observed significantly faster growth rates in the library culture than in the empty vector culture **(Fig. S13)**.

### Yeast Desiccation and Rehydration

Yeast cells grow in distinct phases: lag, log, and stationary. In stationary phase, yeast cells are stressed and compete for resources, so cells prime themselves and show increased viability to desiccation, among other stressors like heat shock. Because we want to profile the extent of protection from the IDP library, we desiccate at mid-log, when they are unprimed. Yeast cells transformed with the library were taken from a glycerol stock and thawed for 30 minutes on ice. A starter culture of approximately 5 × 10^6^ cells/mL was incubated for 5 hours to reach mid-log phase, 1.5 × 10^7^ cells/mL. Cells were then spun down at 6000g for 6 minutes, washed twice with sterile MilliQ H_2_O, and resuspended in one-tenth the volume of water. 300 μL of mid-log cells were aliquoted on filter paper and dried down in a desiccator cabinet containing a saturated salt solution, NaOH, which maintains a relative humidity at 9% ^98^ for two days. Take the remainder of the washed cells for a viability assay and extract the plasmid library to prepare for sequencing. The dried cells were taken from the desiccation cabinet and resuspended in 15 mL of selective media. To ensure dried cells were lifted from the filter paper, the cells and filter paper were incubated in SC-His media in a flask for an hour. Once lifted, a starter culture of 0.2 × 10^7^ cells/mL was inoculated in the turbidostat (purchased from Pioreactor) for outgrowth. The OD600 was calibrated with a cell counter (DeNovix CellDrop FLi), and growth is monitored once every 4 seconds. Yeast cells recovering from desiccation were incubated in 15 mL of SC-His media for approximately 24 hours to keep the samples at mid-log phase before extracting the plasmid library. The turbidostat was programmed to remove 2 mL of cells and to add fresh 2 mL of media to dilute the culture when it reached a density of 2 × 10^7^ cells/mL. Stirring was used in place of shaking at 500rpm, and cells were incubated at 30C.

### Viability by Regrowth Rates

Regrowth rates post-desiccation were obtained from OD monitoring performed in the Pioreactors. A starter inoculation of 0.2 × 10^7^ cells/mL was allowed to recover for 24 hours. Cell density was measured every 4 minutes (**Fig. S13**). Pioreactor settings were as follows: stirring at 500 rpm, 30 °C, removal of 2 mL cells, and addition of fresh 2 mL selective media (SC -His) at cell counts of 2 × 10^7^ cells/mL. OD was calibrated to cell concentration using a cell counter (DeNovix CellDrop FLi). The library was recovered in triplicates, and an empty vector control was used as a baseline recovery post desiccation.

### Illumina sequencing

Plasmids were extracted using the Spin Miniprep Kit (Qiagen) from yeast cells before and after desiccation/rehydration following a modified glass bead lysis protocol. The extracted plasmid library was amplified in 24 cycles and purified with Ampure XP magnetic beads. DNA fragments larger than ∼1 kb were removed using a two-step SPRI bead cleanup (AMPure XP, Beckman Coulter). Briefly, PCR products were incubated with 0.5× bead volume to selectively bind high-molecular-weight fragments, and the supernatant containing smaller fragments was transferred to a new tube. DNA was then recovered from the supernatant by the addition of 1.8× bead volume, followed by two washes with 70% ethanol and elution in 20 μL nuclease-free water. DNA was quantified with a Qubit 2.0 using a dsDNA HS kit. 20 ng of DNA was used for downstream adaptor ligation and barcode amplification with NEBNext Library Preparation kit, following manufacturer’s protocol without size selection. The DNA was quantified on a Qubit fluorometer, and fragment size distribution was checked on an Agilent TapeStation 4200 using the D1000 High Sensitivity ScreenTape assay. Samples were sent out to Azenta for sample preparation and sequencing on their MiSeq platform.

### Illumina Sequencing analysis

For each repeat of the experiment, at least 3 biological samples were sent out for sequencing. Reads were aligned using Bowtie2 and normalized using DESeq2^99^. FastQC files were obtained from Azenta and aligned with Bowtie2^100^ to the reference library. Samtools^101,102^ indexed and sorted the aligned library, and idxstats gave the aligned reads output. DESeq2 differential expression analysis normalized reads between libraries to quantify the fold change between the number of reads before and after desiccation, along with p-values adjusted by Benjamini-Hochberg (BH) correction to control for false discovery rate. The weighted Log2FoldChange average was calculated as 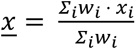 where 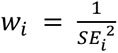 and the pooled standard error was calculated as 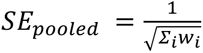.

### Library Sequence Property Analysis

2300 sequences were mapped across 3 repeats. There were a total of 11 template variants, chosen because they had predicted protection ranges from a small preliminary library. At a minimum, 107 variants are generated, up to 282. To plot the correlations between sequence features and variants, a filter of +/− 7% of the WT variant excluded redundant data points to denoise the scatterplot, as many sequences have identical or highly similar sequence feature values to the WT, per the library design. Sequence features were calculated using CIDER^25^, Sparrow^34^, and NARDINI+^59^ (**Table S6)**.

### Yeast Microscopy for IDRs Designed to Form Assemblies

For all yeast microscopy, the base strain was BY4741. Clients and scaffolds were expressed under the TDH3 promoter from a CEN/ARS plasmid containing a URA3 or HIS3 auxotrophic marker. For microscopy experiments, the required strain was streaked onto the appropriate selective plate and incubated at 30°C to obtain single colonies. The day before imaging, a single colony for each needed strain was selected and grown in 6-well plates in 2mL liquid cultures at 30°C 185 RPM. The next morning ∼3 hours prior to imaging, each strain was subcultured at a 1:1 ratio to maintain log phase growth. One hour prior to imaging, slides were prepped. Imaging utilized Ibidi USA µ-Slide 18 Well, Glass Bottom - #1.5H Glass Bottom Coverslip (Ibidi USA Cat. No. 81817, Fisher Scientific Cat. No. NC1752257). The 18-well µ-Slide was treated with 40 µL of 1 mg/mL Concanavalin A (Thermo Scientific Chemicals, Cat. No. J61221-MD) for 15 minutes to help adhere yeast to the slide to keep them stationary during imaging. Next, 1mL of each culture was spun down and resuspended in 100 µL of the remaining supernatant. At this point, the Concanavalin A solution was aspirated from each well. Each well was then washed with 40 µL of water. Next, the water was aspirated, and 40 µL of culture was added to the well and left to incubate for five minutes. Next, the non-adherent cells were aspirated off, and 40 µL of fresh media was added to the well. At this point, the yeast were immediately taken for imaging. Imaging was carried out at the Washington University in St. Louis Center for Cellular Imaging (WUCCI) using the Zeiss LSM 880 Confocal with Airyscan microscope equipped with a Plan-Apochromat 100x/1.40 NA Oil DIC M27 objective. The mTagBFP2 imaging used a 405 nm diode laser at 1%, with the PMT detector gain set to 720, a detection wavelength of 410-530 nm, and ∼1.0 Airy Units. The mScarlet3 imaging used a 561nm DPSS laser at 0.75%, a PMT detector gain of 720, a detection wavelength of 582-754 nm, and ∼0.7 Airy Units. The mTurquoise2 imaging used the 458 nm Argon laser at 1.0%, PMT detector gain set to 750, detection wavelength from 462-580nM, and ∼0.99 Airy Units.

### Quantifying partition coefficients for colocalization analysis

Quantification of partition coefficients for colocalization analysis of the GOOSE-designed IDRs with the synthetic self-assemblies was done using a custom Python script using scikit-image and SciPy libraries. The cells were imaged as z-stacks, and a substantial number of out-of-focus planes were in the data. To avoid data accuracy issues caused by out-of-focus cells, the first and last z-planes of the z-stack were discarded, leaving only the middle 50% for analysis. First, cells were segmented using the client channel, slice by slice. The global background was then determined using the median absolute deviation of the image array. The foreground was segmented using Li’s Minimum Cross Entropy algorithm, which thresholds images with heavily skewed object populations (such as self-assemblies)^103^. Binary masks were then further refined using median filtering, morphological closing, and hole-filling. Touching cells were separated using a Euclidean distance transform coupled with a watershed algorithm, and then further filtered by size exclusion to remove noise and truncated cells. Finally, the cell boundary was uniformly shrunk by 10 pixels to avoid including the area outside the cell. This was necessary to avoid skewing the average intensity calculated for the client signal that did not overlap with the assemblies (this would artificially reduce the average client signal and thus artificially increase calculated partition coefficients). Next, assemblies were segmented independently using the scaffold channel. This used a two-pass statistical approach: first, the cellular foreground was identified using a 50x the median absolute deviation threshold, and then refined using median filtering. This was followed by morphological opening to remove noise, hole filling, and separation of individual assemblies using a local-maxima watershed. Assemblies with fewer than 50 pixels or with intensity values below 2,000 intensity units (values empirically determined) were then filtered out.

Once the cell boundaries and assembly boundaries were identified for every z-plane, a “2.5D axial linking approach” was used. In this approach, assembly masks generated on individual 2D slices were evaluated for overlap across the z-axis. Overlapping assemblies were assigned a single identifier (assuming that assemblies in the same location across multiple z-planes were the same). For assemblies spanning multiple z-planes, each plane was examined, and the plane with the highest intensity was used to assess colocalization of the client with that assembly. All other z-planes were discarded, such that each assembly was only quantified once. The average fluorescence intensity was measured for the client protein inside the assembly mask (I_in_) and in the surrounding area (I_out_). In addition, the background fluorescence (I_bg_) was estimated from non-cell regions and subtracted from the total fluorescence (I). The partition coefficient was calculated as 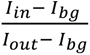. For quantification, Li’s Minimum Cross Entropy threshold was applied to eliminate cells that were expressing the client at too low a level. As a final quality control step, we set up our analysis to output .png file showing the raw channels for the assembly and the client, next to what was used as the mask for ‘inside the assembly’ and ‘outside the assembly’, such that we could examine each data point individually and ensure there were no abnormalities with the quantification. For example, if an assembly was localized to the nucleus for one of the constructs that tended to have strong nuclear client signal, that data point would be excluded because it would measure the overlap in assembly signal to strong client nuclear signal, which would suggest strong colocalization signal that is largely due to the nuclear signal of the client rather than true colocalization with the scaffold.

To assess whether individual constructs differ significantly in their partitioning behavior, we performed pairwise Mann-Whitney U tests (two-sided) on all unique pairs of constructs within each scaffold condition. The Mann-Whitney U test was chosen as a non-parametric alternative that makes no assumptions about the underlying distribution of partition coefficients. For each scaffold, given *k* constructs, we computed 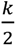 pairwise comparisons. To control the family-wise error rate across the multiple comparisons within each scaffold, p-values were corrected using the Holm-Bonferroni step-down procedure, which provides strong control of the family-wise error rate while being uniformly more powerful than the classical Bonferroni correction. Corrected p-values were considered significant at α = 0.05, with additional thresholds at 0.01 and 0.001 for graphical annotation. All statistical tests were performed using SciPy (**`scipy.stats.mannwhitneyu`**), and multiple testing correction was applied via statsmodels (**`statsmodels.stats.multitest.multipletests`**, method “holm”)^104^.

### Coarse-grained simulations

Coarse-grained molecular dynamics simulations were used to assess how well GOOSE-generated sequences conform to target ensemble dimensions. Simulations were performed using the LAMMPS simulation engine^105^ using a modified version of the Mpipi^89^ parameters, Mpipi-GG^45^. Starting positions for IDRs were generated by assembling beads as a random coil in the excluded volume limit (i.e., where beads do not overlap). From this position, an energy minimization protocol was carried out with a maximum of 1,000 iterations. Simulations were then carried out with an implicit salt concentration of 150 mM and a temperature of 300 K. Simulation analysis was performed using MDTraj^106^ and SOURSOP^107^. For simulations of IDR designed to match specific radii of gyration or end-to-end distances (200 residues, Fig. **S7**), all sequences were run in triplicate for 200,000,000 steps with a 20 fs timestep for a total of 12 µs per sequence. The first 1,000,000 steps for each simulation were discarded as equilibration. After equilibration, output coordinate positions for each trajectory were recorded every 100,000 steps, for a total of 1,990 steps per simulation. This simulation length was chosen based on prior work to benchmark appropriate simulation lengths to obtain robust conformational sampling^34^. Error bars (shown in **Fig. S7**) show that the variability between independent replicas is minimal (on the order of the marker size in the figure), again confirming that simulations sufficiently sample the conformational landscape. The simulation used a 500 Å^3^ box with periodic boundary conditions.

### Data Availability

GOOSE source code is available at https://github.com/idptools/goose/

GOOSE documentation is available at https://goose.readthedocs.io/

Data and analysis scripts used for figures and analysis in this paper are available at https://github.com/sukeniklab/GOOSE_2026 and at https://github.com/holehouse-lab/supportingdata/tree/master/2026/GOOSE_2026

Raw sequencing data are deposited at Zenodo and are publicly available at https://doi.org/10.5281/zenodo.18717145

## ACKNOWLEDGEMENTS

This work was funded by a US National Science Foundation (NSF) IntBio Collaborative Research Proposal to S.S. (#2128067) and A.S.H (#2128068), an NSF BII Award (Water and Life Interface Institute, WALII, NSF DBI grant #2213983) to S.S. and A.S.H., an NSF CAREER award (#2338129) to A.S.H., a Research Grant from the Human Frontiers in Science Program (HFSP) to A.S.H. (RGP0015/ 2022), and by the US NIH under award R35GM137926 to SS. KG was supported by a fellowship from NSF-CREST Center for CCBM at UC Merced, Grant No. NSF-HRD-1547848. K.H. was supported by the US NIH under award number T32GM141862. TB was supported in part by funds from the Syracuse University School of Arts and Sciences. J.M.L. was supported by an NSF GRFP (DGE-2139839). We also thank members of the Water and Life Interface Institute (WALII) for helpful discussions. The authors thank Max Staller, Marcin Suskiewicz, Susan Dutcher, Tim Lohman, Gaya Amarasinghe, Rohit Pappu, Magnus Kjaergaard, Stephen Plassmeyer, Jackie Pelham, Garrett “mother vat” Ginell, and FNZ for many useful discussions and comments on the manuscript. A.S.H. is a member of the Scientific Advisory Board for Prose Foods. All other authors declare no conflicts of interest.

