## Supplementary Information for "Rational design of disordered proteins for systematic sequence-to-function investigation"

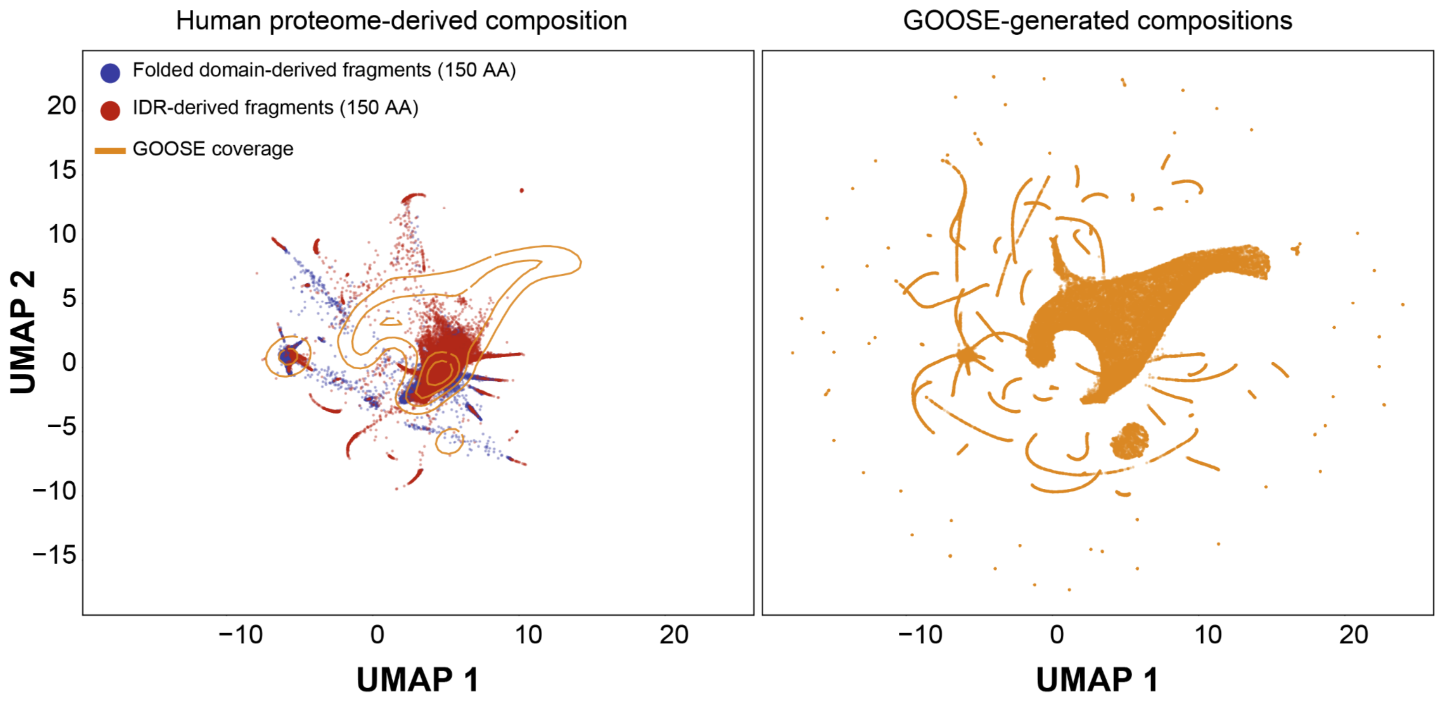


**Fig. S1. Comparison of Natural and Synthetic Sequence Space via UMAP**
Uniform Manifold Projection (UMAP) showing the projected compositional space occupied by 150-amino acid fragments taken from the human proteome (left) or generated by GOOSE (right). UMAP was generated by embedding sequences into an L=20 vector, where each element corresponds to the fraction of each amino acid. Human proteome fragments were identified by segmenting the human proteome into disordered regions and folded domains, then selecting each 150-amino-acid subfragment within those regions with a step size of 50 amino acids. This yielded 33,626 IDR fragments and 85,098 folded fragments. For GOOSE, we generated a minimum of 10 sequences by titrating across amino acid fractions, GOOSE properties (NCPR, FCR, hydropathy, kappa), and amino acid classes (aromatic, aliphatic, polar, positive, negative, glycine, proline, cysteine, histidine), titrating from minimum to maximum possible values in a stepsize of 0.1 across all parameters described. Overall, this yielded 67,427 150-residue GOOSE-derived segments, generated in ~2.5 hours. We note that the generation time can be reduced to minutes by avoiding the extremes of possible sequence composition. GOOSE-generated sequences encompass almost all of the IDR-centric regions, yet substantially expand beyond natural sequence space (compare orange contours on left with natural sequence space in red).

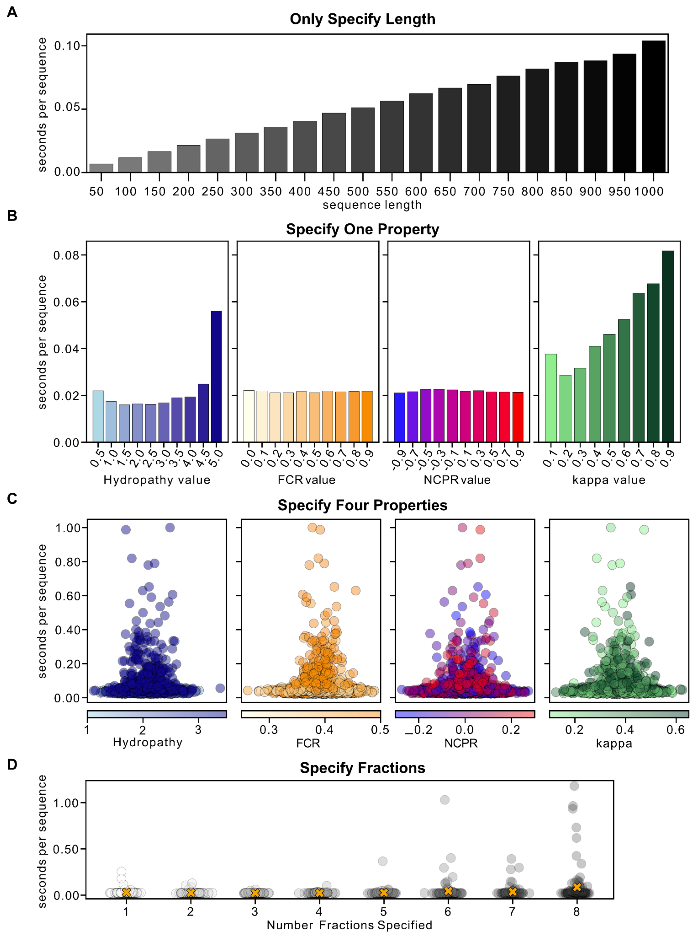


### Fig. S2. GOOSE performance assessment

GOOSE can rapidly generate sequences with various specified properties. **(A)** Time to generate a sequence where only the length is specified. Times are average values from 200 generated sequences. (**B)** Time to generate sequences where only a single property is specified. All sequences are 200 amino acids in length. Times are average values from 200 generated sequences. **(C)** Time to generate sequences where four properties are specified. Depending on the combination of sequence properties, GOOSE can take longer to generate a sequence. Each point represents the duration required to generate an individual sequence. Each plot shows the property for that sequence. All sequences are 200 amino acids in length. A total of 1,656 generated sequences are shown titrating across hydrophobicity, FCR, NCPR, and kappa values. **(D)** Time to generate sequences where different numbers of amino acid fractions are shown. A total of 100 sequences were generated for each number of fractions specified. Each point is an individually generated sequence. The orange 'X' shows the average value across the 100 sequences. For each sequence, the amino acid and its corresponding fraction were randomly chosen within a range constrained by the remaining fraction available to specify (fraction cannot go over 1) and the maximum fraction of that amino acid that can be specified while retaining sequence disorder. Maximum fractions were determined by attempting to generate a sequence of 100 amino acids in length at each fraction for every amino acid between the decimal fraction values of 0.01 to 1.00. For each fraction value, the sequence was populated with the necessary number of amino acids of interest, and then the rest of the sequence was generated by populating the sequence with any amino acid other than the amino acid that had its maximum fraction determined. 500,000 sequences were attempted at each fractional value and then checked to be disordered using metapredict V2 with a cutoff of 0.5.

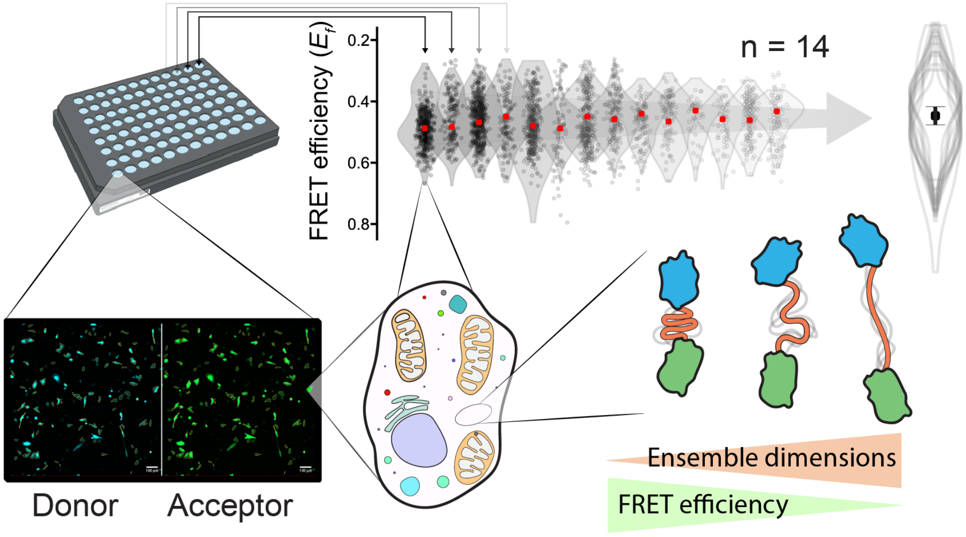


**Fig. S3. Overview of in-cell FRET approach**

An in-cell fluorescence reporter assay combined with a library of rationally designed sequences enables us to determine how IDR sequence properties dictate ensemble properties in the cellular environment (bottom right). Experiments are performed on a 96-well imaging plate (top left), where each well is individually transfected with plasmids encoding for the FRET construct. Wells are counted as individual biological repeats if they include 30 or more cells expressing the construct. Each cell is analyzed to calculate FRET efficiency ($E_f$) from donor and acceptor fluorescence (bottom left). Individual cells are collected to create a violin plot for each well, and this is done across multiple independent measurements (14 in this case). The median of each violin is averaged, and the standard deviation between these medians is reported to provide a highly reproducible assessment of sequence-specific FRET efficiency (top right). Only the overlayed violin plots shown on the top right are reported in all other figures; the average median is shown as a circle, and the SD of medians as error bars.

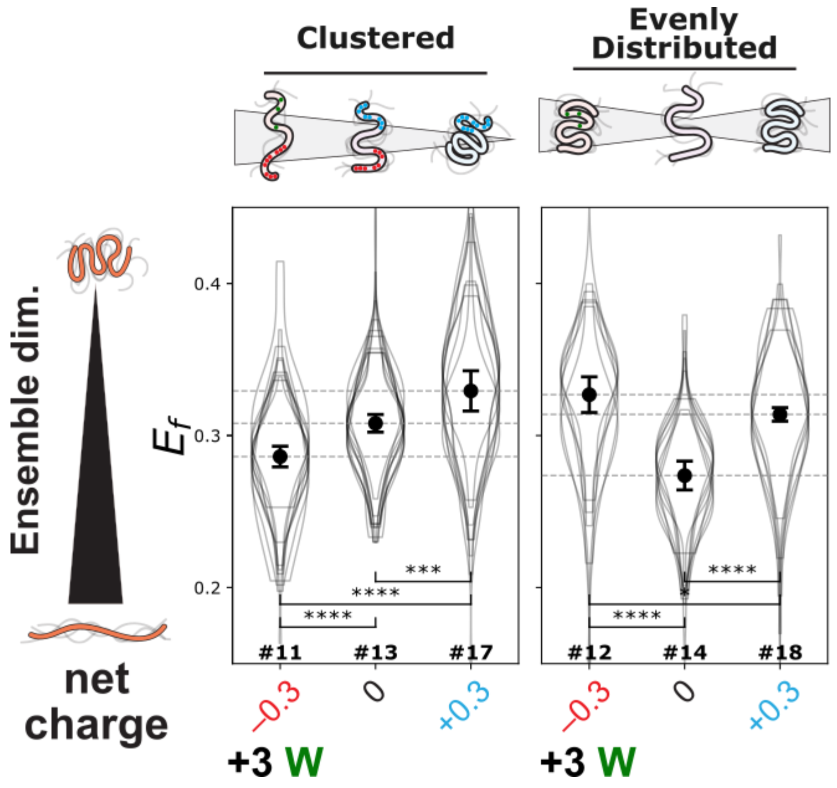


**Fig. S4. Interplay of aromatic and charged residues**

$E_f$ for sequences with a constant FCR of 0.3 and a variable net charge (-0.3, 0, +0.3), with the net negatively charged sequences having 3 aromatic residues (tryptophans) introduced. When charges are clustered at one end of the IDR and tryptophans at the other, the sequence shows behavior (left) comparable to that of other variable charge triplets (**Fig. 2G**). However, when charges are evenly distributed, addition of tryptophans leads to significant ensemble compaction (right).

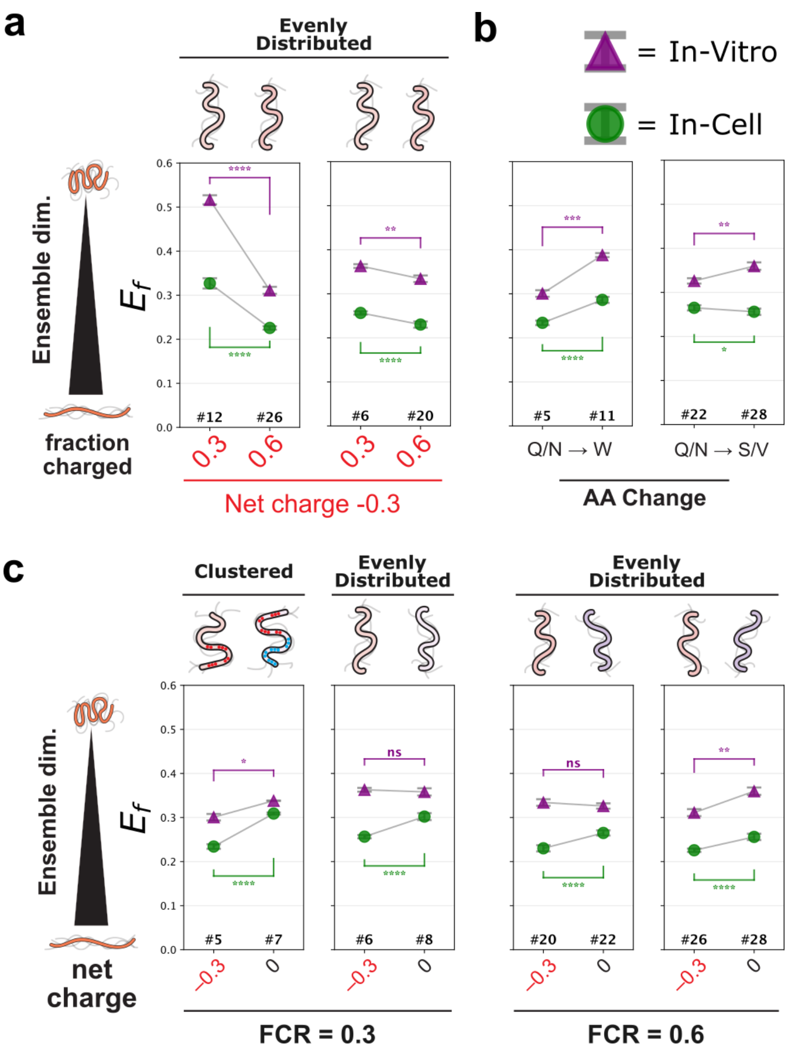


**Fig. S5. *In vitro* vs. in cell comparison of disordered protein ensembles.**

Pairwise $E_f$ comparisons of *in vitro* (purple) and in-cell (green) data. *In vitro* data points are the mean of biological replicates (n = 3), and error bars are their standard deviation. In cell markers and error bars are the average and standard deviation of the medians of all biological repeats (n > 5) (Fig. 2). Significance is determined by a Welch's t-test. **(a)** $E_f$ for sequences with evenly distributed charges and a constant net charge of -0.3. **(b)** $E_f$ for sequences that introduce aromatics and modulate hydrophobicity. In both comparisons, ensemble compaction is expected: overall, removing Q/N promotes it due to the loss of hydrogen-bonding participants. The addition of hydrophobic aromatics (W) that also engage in self-attractive pi-pi and cation-pi interactions strengthens this effect. Mutation to relatively more hydrophobic residues, S and V, would also be expected to contribute to ensemble compaction. **(c)** $E_f$ comparisons for sequences that modulate FCR, net charge, and charge clustering. In-cell structural ensembles compact monotonically when positive charges are added, with charge patterning and FCR having little effect on this relationship. *In vitro* constructs behave in a more variable, sequence-dependent manner.

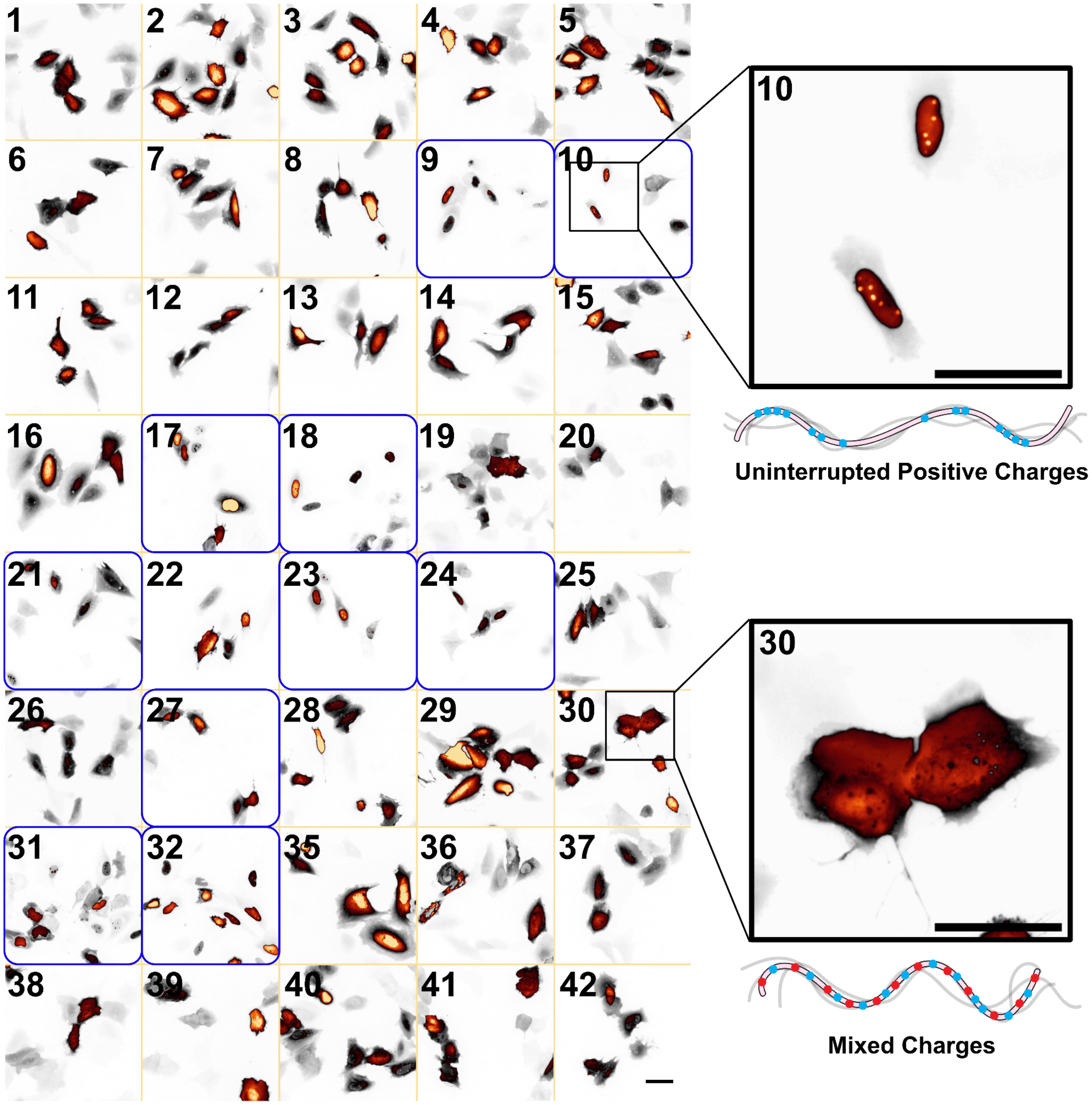


**Fig. S6. Cellular localization and puncta formation of GOOSE-generated sequences**

Representative images of the direct acceptor channel from all 40 constructs generated by GOOSE and used for $E_f$ measurements. Numbers in the top-left of each box correspond to sequence numbers from **Tables S1 and S2**. Blue boxes around sequences indicate sequences with two or more patches of three or more positive amino acids. Almost all constructs with runs of positively charged residues (excluding #27, 9/10) display nuclear puncta consistent with inclusion into RNA-ribonucleoprotein bodies, as well as strong nuclear localization (e.g., #10). When sequences lack clustered positive charges or contain mixed positive and negative charges, they display a diffuse signal (e.g., #30). Scale bar = 50 μm.

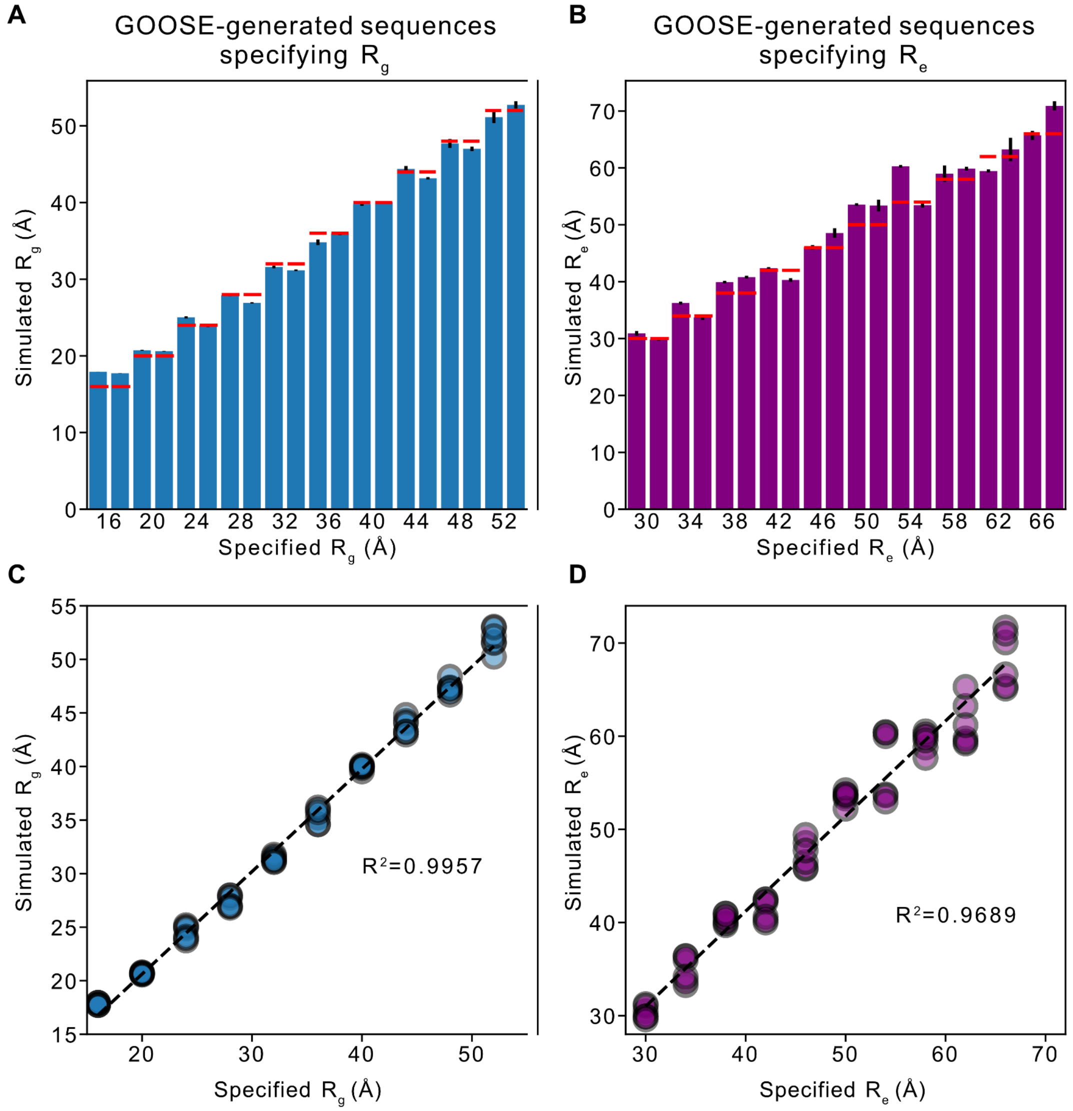


**Fig. S7. Generation of IDRs with desired ensemble properties**

All sequences generated were 200 amino acids in length. **(A, C)** For $R_g$ , two sequences with dimensions between 16 Å and 52 Å at intervals of 4Å (20 sequences total) were generated. **(B, D)** A similar approach was used to specify $R_e$, except a range of 30 Å to 66 Å was used. After sequence generation, coarse-grained molecular dynamics simulations were run as described in the *Methods*. For bar plots (A, C), bars are equal to the mean of the average $R_g$ or $R_e$ of the triplicate for each sequence, error bars are the standard deviation between the means for each triplicate, and the x-axis labels denote the $R_g$ or $R_e$ specified for each sequence (two sequences per specified dimension). Red lines indicate the specified $R_g$ or $R_e$ during sequence generation. Each point in the scatter plots (B, D) shows the average dimension for each simulation triplicate, for both sequences, at the desired $R_g$ or $R_e$ value (y-axis), with the specified dimension during sequence generation on the x-axis. The $R^2$ values were calculated using the mean value of the triplicate for each sequence vs. the specified $R_g$ or $R_e$ during sequence generation for each sequence. $R^2$ reports the coefficient of determination (Pearson's r, squared).

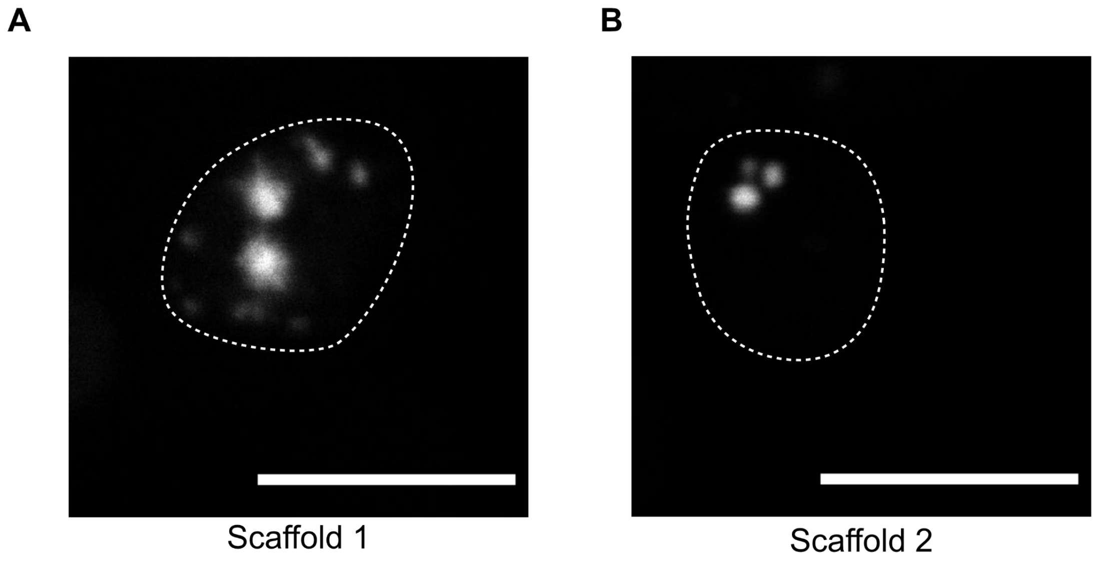


**Fig. S8. Scaffold-only constructs assemble in the absence of client**

Image of yeast expressing scaffold constructs. **(A)** mTagBFP2-Scaffold1 and **(B)** mScarlet3-Scaffold2 alone. Both scaffolds were expressed under the TDH3 promoter. Cell outlines are shown as dashed lines. Only cells with visible expression show outlines. The scale bar is 5μm.

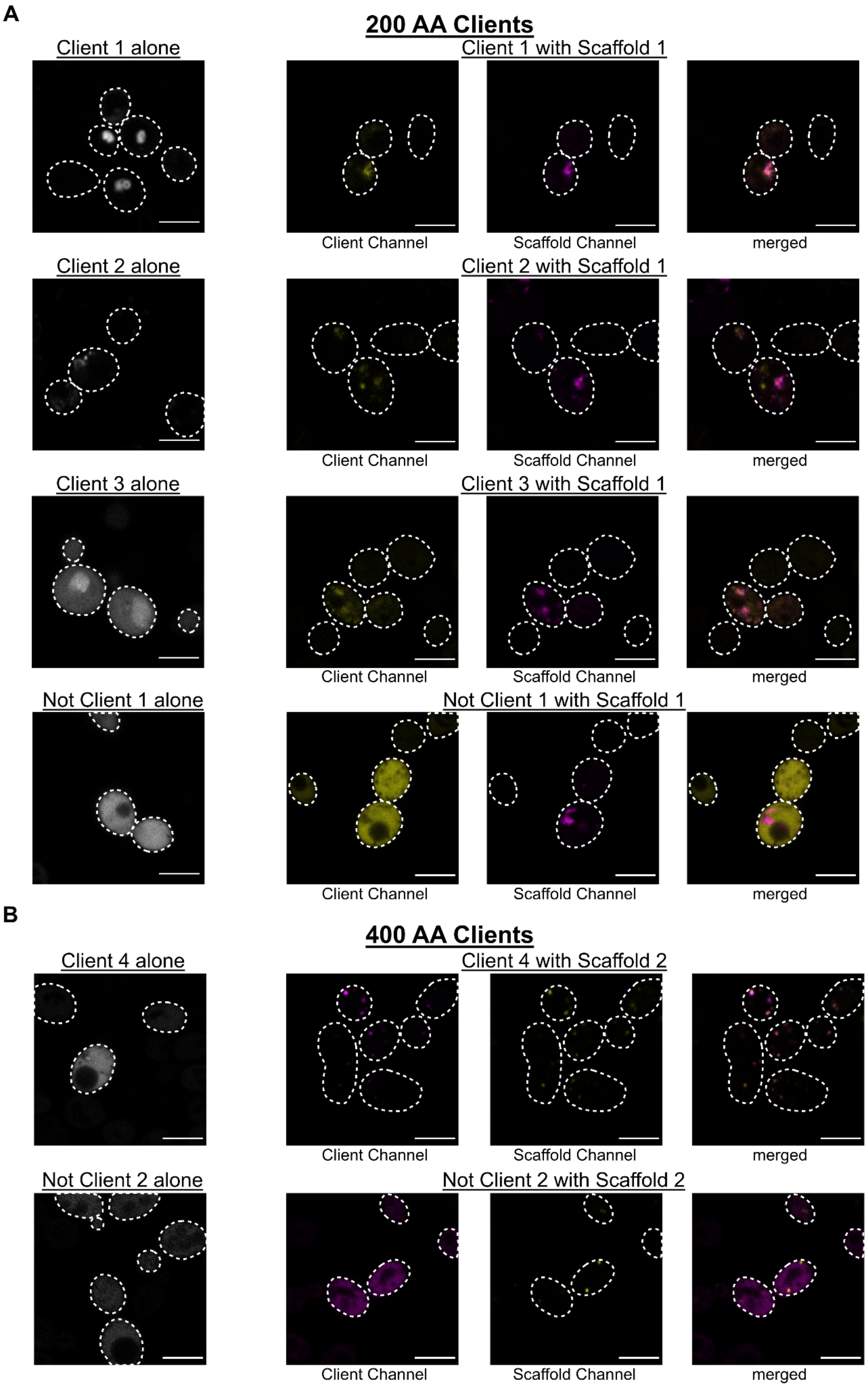


**Fig. S9. Scaffold-dependent recruitment of synthetic client constructs in yeast**

Image of yeast expressing client construct alone (furthest left) or client construct with scaffold (right three columns). All scaffolds and clients were expressed under the TDH3 promoter. The scale bar is 5μm for all images. **(A)** The scaffold construct was mTagBFP2-Scaffold1. All clients were 200 amino acids and used a mScarlet3 fluorophore. **(B)** The scaffold construct was mScarlet3-Scaffold2. All clients were 400 amino acids in length and used the mTagBFP2 fluorophore.

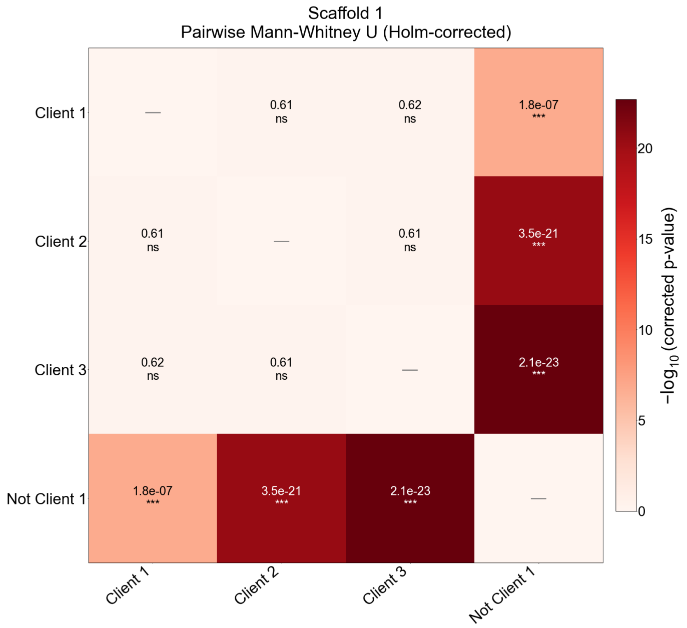


**Fig. S10. Statistical comparison of client recruitment across scaffold conditions (Scaffold 1)**
Mann-Whitney U tests (two-sided) with Holm-Bonferroni correction on all unique pairs of constructs within each scaffold condition as described in **Fig. 4H**.

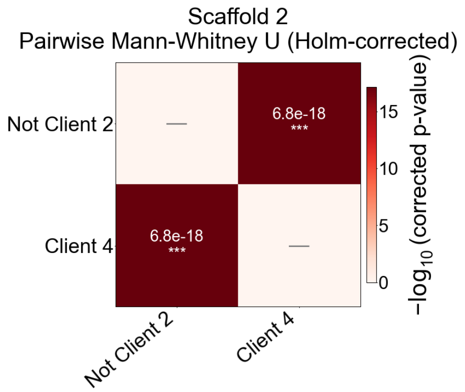


**Fig. S11. Statistical comparison of client recruitment across scaffold conditions (Scaffold 2)**
Mann-Whitney U tests (two-sided) with Holm-Bonferroni correction on all unique pairs of constructs within each scaffold condition as described in **Fig. 4I**.

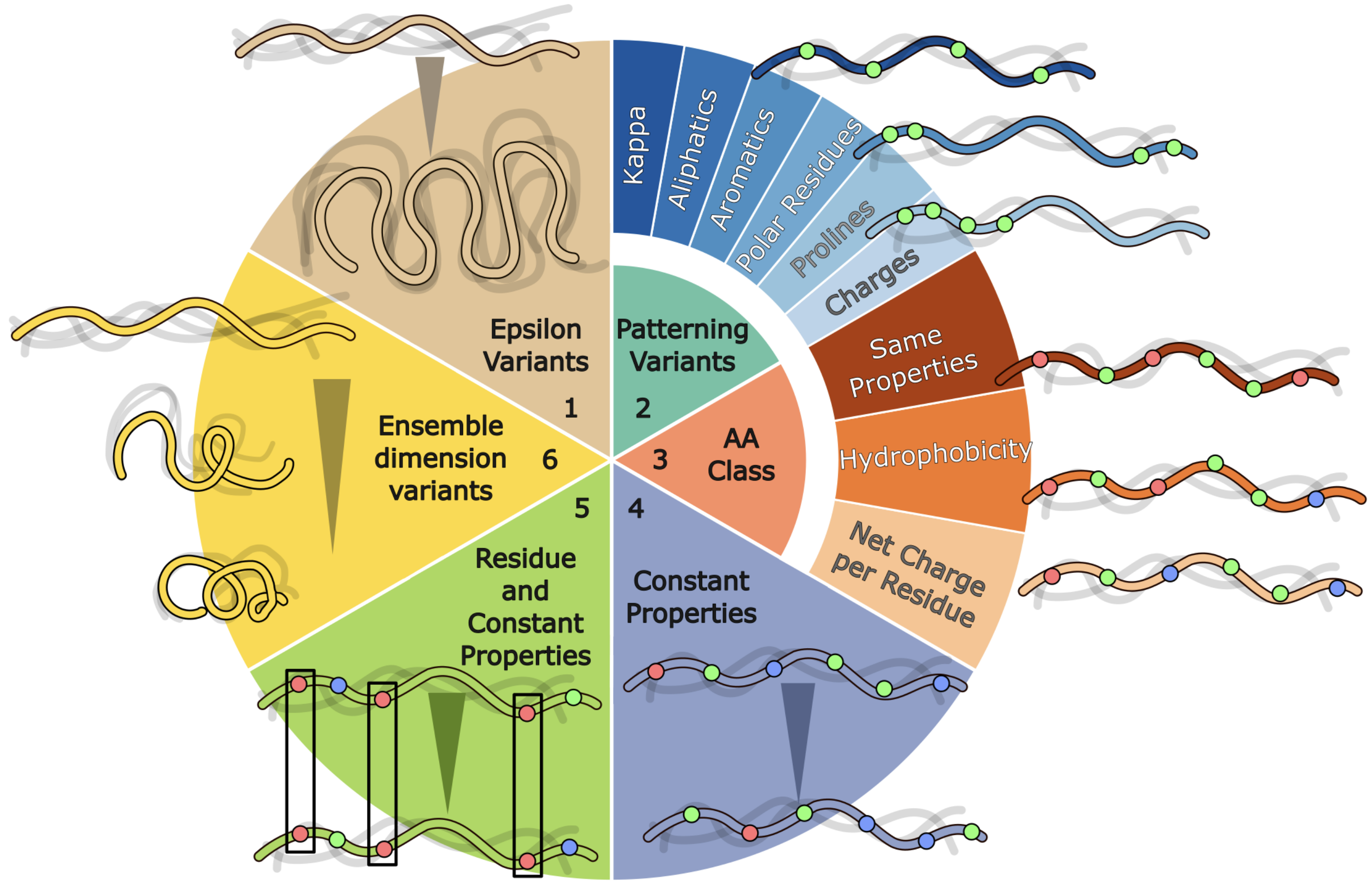


**Fig. S12. Design strategies for the desiccation protection sequence library**

A schematic of all variation types in the desiccation protection library. Variants were organized into six major categories based on the level of sequence constraint applied. The first category, epsilon variants, increases or decreases the predicted self-attraction/repulsion as predicted by FINCHES. The second category, patterning variants, keeps the amino acid sequence itself unchanged and instead explores sequence space through positional rearrangement. This category includes variants that increase or decrease charge clustering $\kappa$, as well as variants that modulate the clustering of specific residue groups, including aliphatics, aromatics, polar residues, histidine (when present), or prolines (when present). The third category, constant amino acid by class, holds the number of residues per amino acid class fixed and includes three subtypes: (1) variants where overall chemical properties are preserved through substitutions restricted to chemically similar residues (e.g., Val and Ile, Ser and Thr); (2) variants that increase or decrease hydrophobicity while keeping the amino acid class order constant; and (3) variants that increase or decrease the net charge per residue (NCPR). The fourth category, constant properties variants, constrains only the overall chemical properties while allowing the number of residues per amino acid class to vary freely. The fifth category, residue and constant properties, holds a single residue class and overall properties constant while allowing all other aspects of the sequence to vary freely. The sixth category, ensemble dimension variants, alters the end-to-end distance ($R_e$) by changing the spatial distance between the N- and C-termini without altering sequence composition. Together, this constrained exploration of sequence space enables downstream analysis and hypothesis generation that would otherwise be inaccessible through single-amino-acid mutations alone.

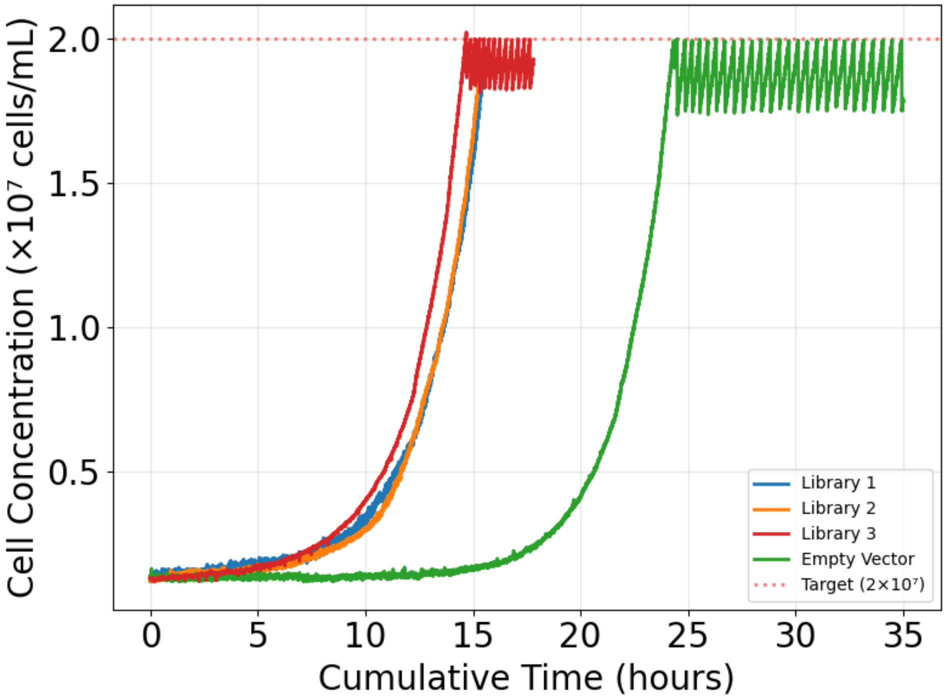


**Fig. S13. Growth kinetics and automated culture maintenance in pioreactors**

Plotted cell concentrations of libraries (blue, yellow, red) and an empty vector control (green) over time, measured by optical density in Pioreactors. The optical density was calibrated and converted to cell concentration (cells/mL) by correlating OD with cell counts (DeNovix Celldrop FLi). The Pioreactor was programmed to remove 2 mL of cells and add 2 mL of fresh media to dilute the culture when the cell concentration reached $2.0 \times 10^7$ cells/mL. This ensures the culture remains in log phase. Removal and addition of fresh media is visualized in library 3 (red) at 15 hours, and in the empty vector (EV) control at 25 hours.

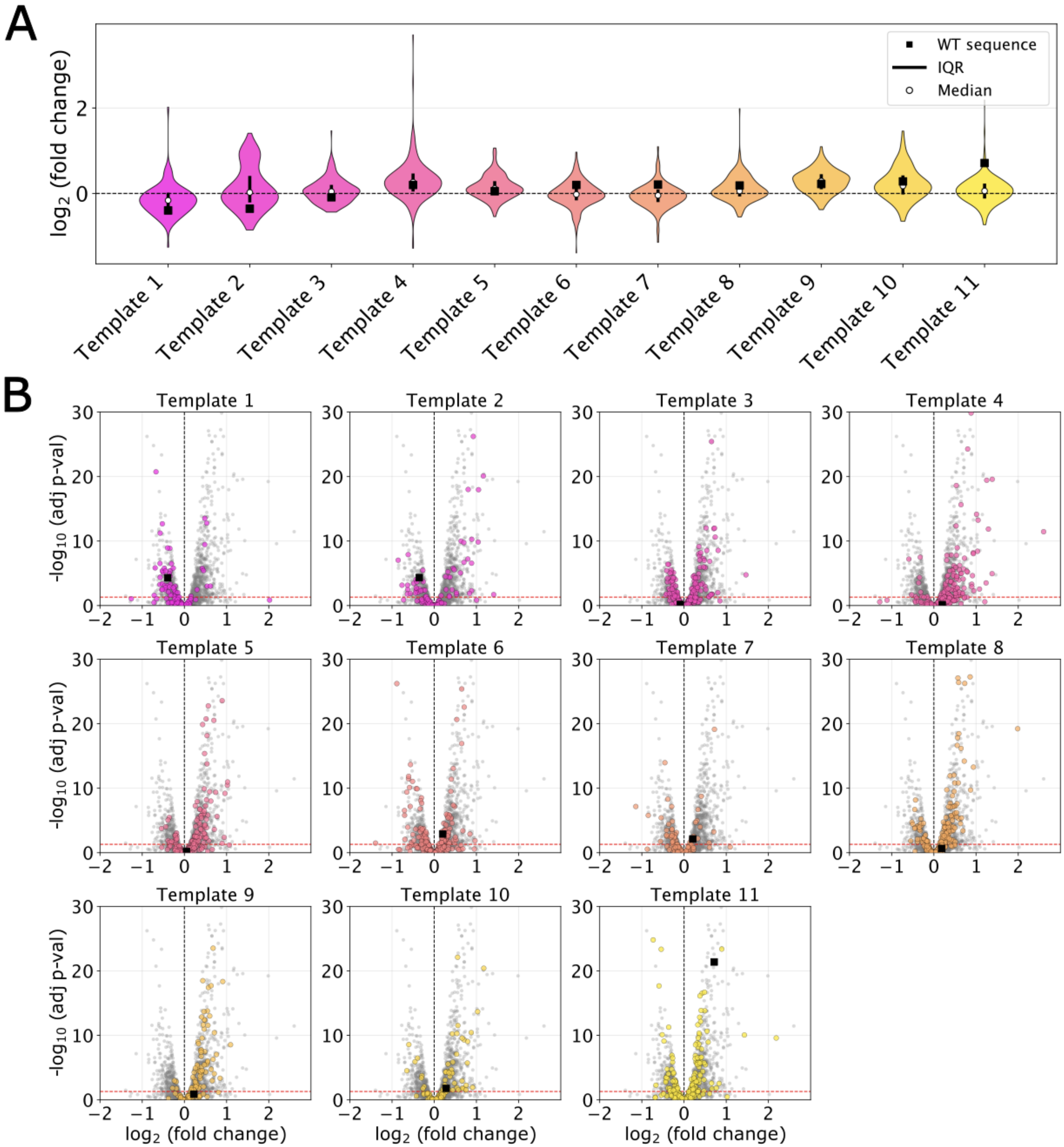


**Fig. S14. Distribution and significance of log2-fold change across sequence variants**

(**A**) Violin plots of all templates and their corresponding weighted log2FoldChange values across 3 separate repeats of the experiment. The wider the violin, the more frequently that specific log2FoldChange value occurs. Visualized were the WT template value, the IQR, and the median (white circle). (**B**) Volcano plots of the calculated log2FoldChange and the -log10 (adjusted p-value) highlight the original template sequence (black square) and its respective variants generated by GOOSE.

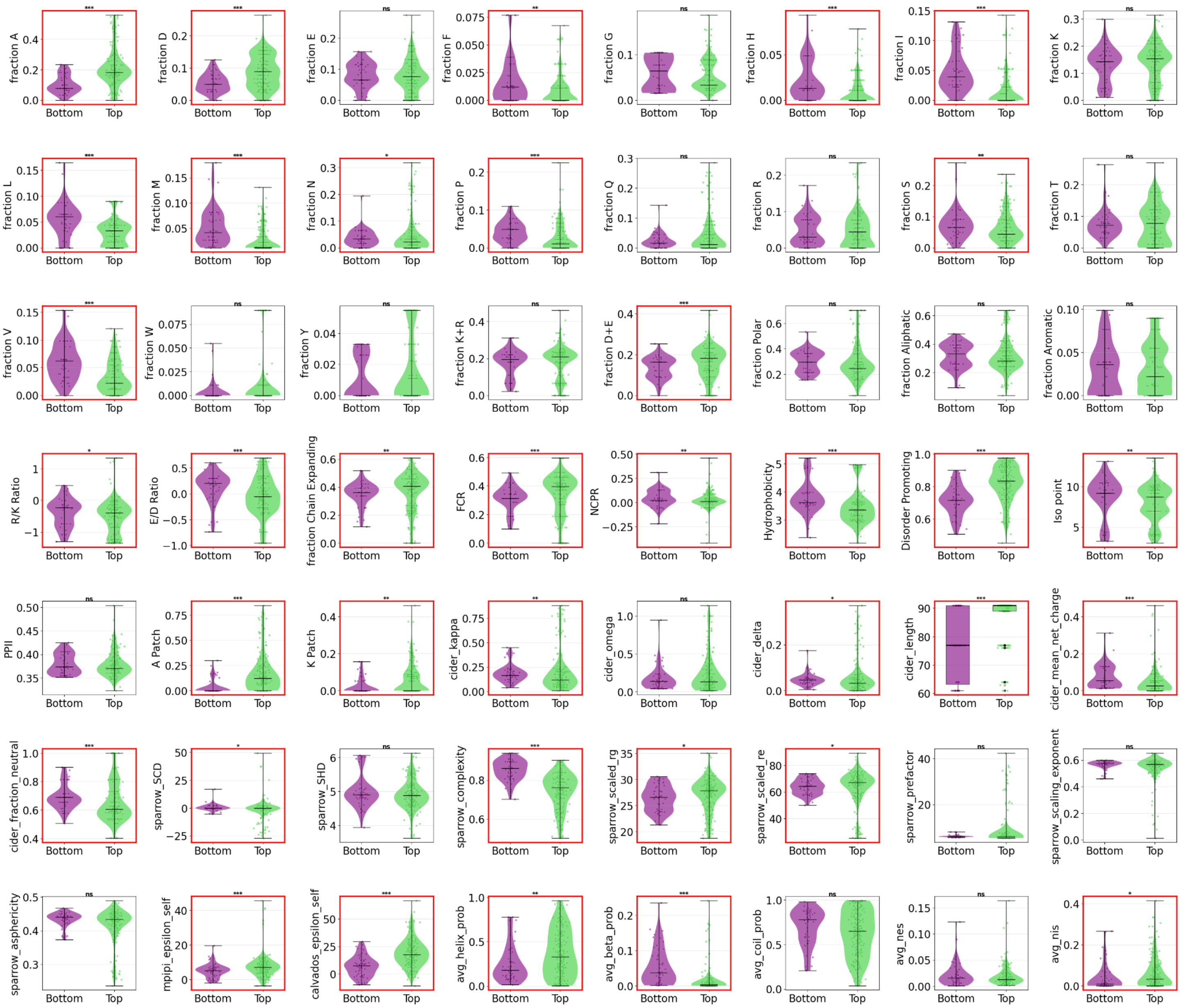


**Fig. S15. Comparative distribution of top and bottom-performing sequence variants**

Plotted violin plots of the worst performing sequences (labeled ‘bottom’ and colored purple: determined by a log2FoldChange of -0.5 or less, statistically significant), and top performing sequences (labeled ‘top’ and colored green: determined by a log2FoldChange of 0.5 or greater, statistically significant). Significantly different distributions were determined by a two tailed Mann Whitney U-test, corrected for multiple testing using the Benjamini-Hochberg procedure to control the false discovery rate (FDR), and are shown with a red border.

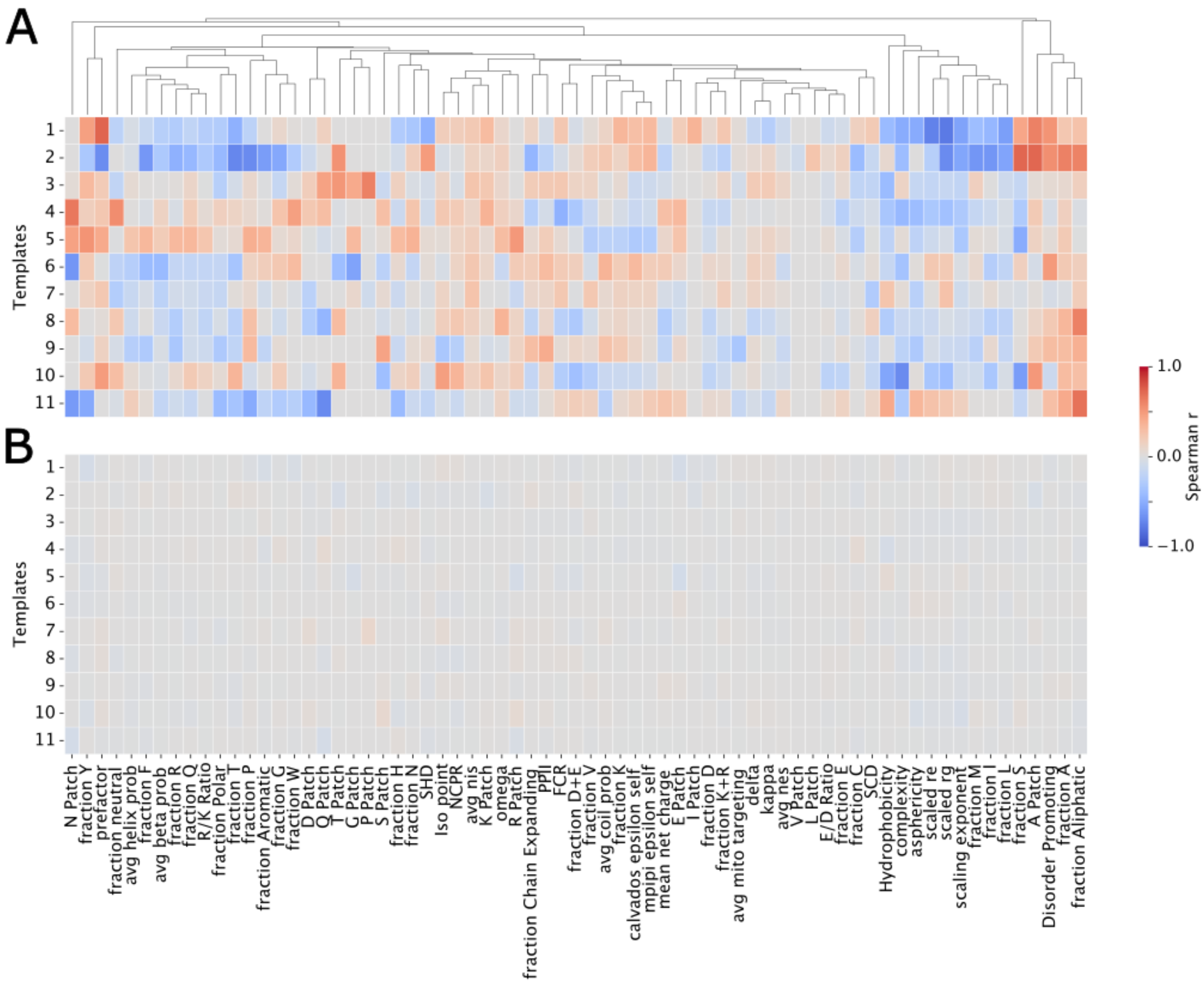

**Fig. S16. Permutation analysis to validate experimental parameter correlations**

To determine the likelihood of false positives for strong correlations, the average fold changes and their respective errors were randomly shuffled, and the Monte Carlo Spearman's r was recalculated for every shuffle. This was calculated for 100 permutations, then the correlations were averaged per parameter. Experimental correlations (**A**) were compared to randomly shuffled correlations **(B).** These permutations revealed that none of the parameters showed correlation after shuffling. Explanations on each of the parameters are given in **Table S6**. Parameters were removed if they were redundant or if, after filtering within +/-7% of the WT parameter value, n < 10.

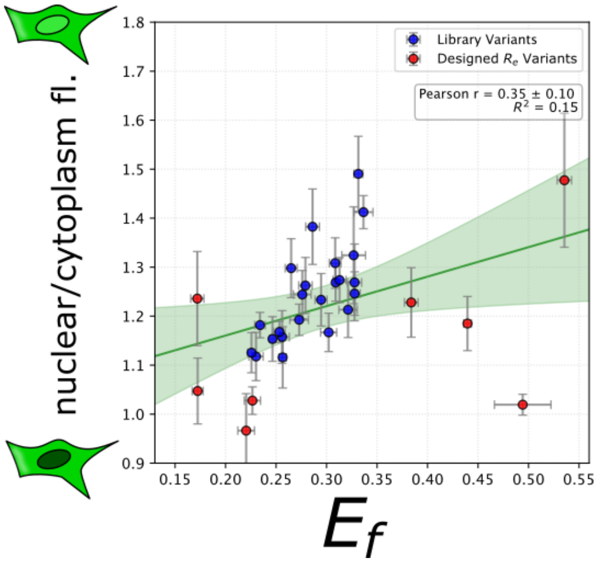


**Fig. S17. Correlation between ensemble dimensions and passive nuclear localization**

Correlation between constructs' ensemble dimension ($E_f$) and nuclear localization (nuclear/cytoplasm fl.) with designed $R_e$ sequences included (red). Sequences that displayed localization consistent with the presence of an NLS signal (**Fig. S6**) were excluded from this graph, as those do not inform on the relationship tested here, namely passive diffusion through the nuclear pore complex. Designed $R_e$ sequences poorly recapitulate the library trend, possibly due to a deviation from molecular grammars present in the original library. For example, sequences # 41 and 42 (far left points) have NCPR = ~-0.66, while # 35 and 36 (far right points) have aromatic residue fractions of ~0.33.

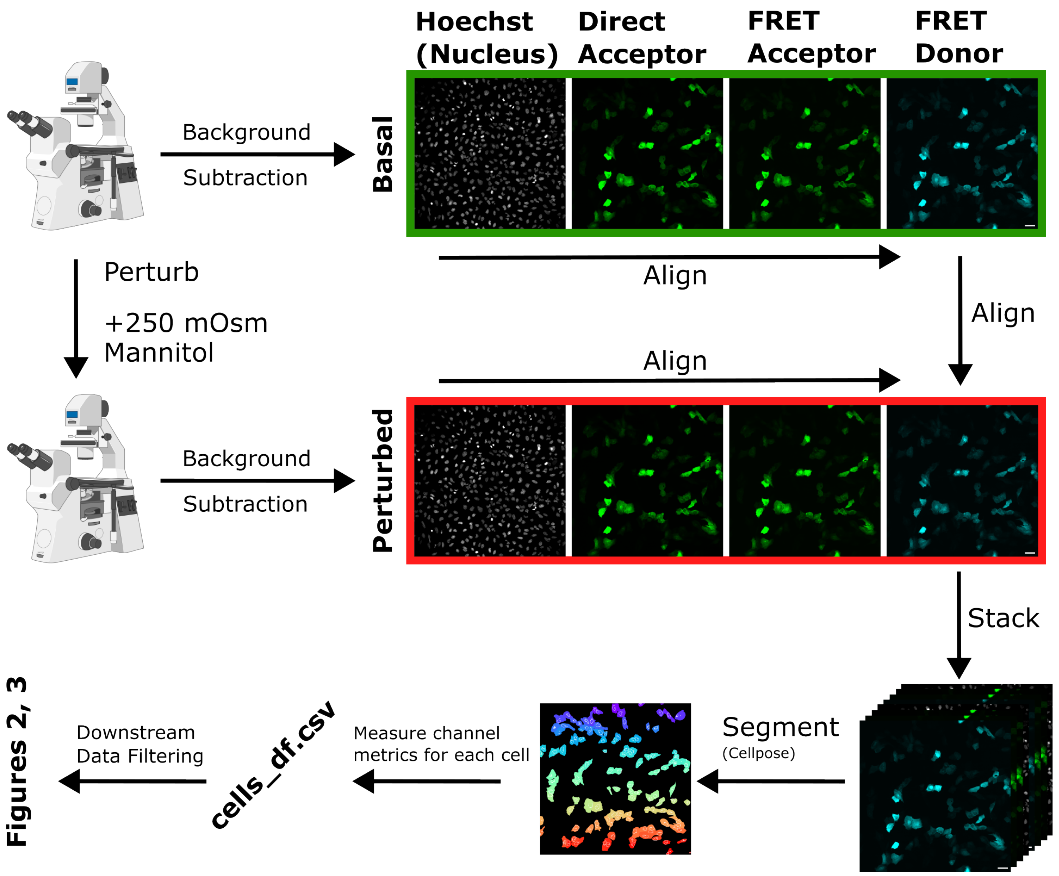


**Fig. S18. Image processing pipeline for quantifying $E_f$, cell compaction, and localization**

Schematic showing the analysis pipeline from the initial collection of images to per-cell dataframe creation. Briefly, images are collected before and after osmotic perturbation, then stacked and aligned, and then segmented to create masks for each cell. Fluorescence intensity metrics per cell are collected, and $E_f$ and localization metrics (directA_N/C)are calculated. See methods (**image analysis**) for a full description. Scale bar = 50 μm.

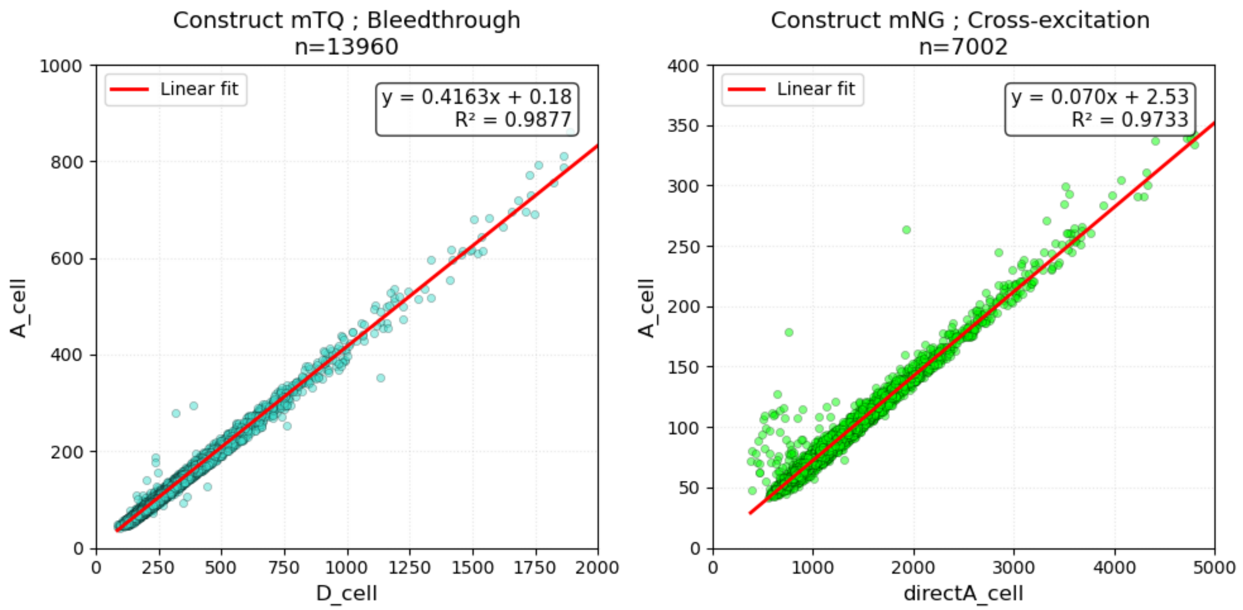


**Fig. S19. Linear regression and correction of fluorescence artifacts**

Linear regressions are used to calculate fluorescence artifact correction values for the acceptor channel. Bleedthrough of uncoupled mTurquoise fluorophores into the acceptor channel when excited (left). Cross-excitation of uncoupled mNeonGreen fluorophores when excited by FRET excitation light (right). Corrected acceptor is then calculated from the slope of these regressions, per cell, in FRET constructs, by the following equation: $Acceptor_{corr} = Acceptor - (.070 \times direct\ Acceptor) - (0.4163 \times donor)$

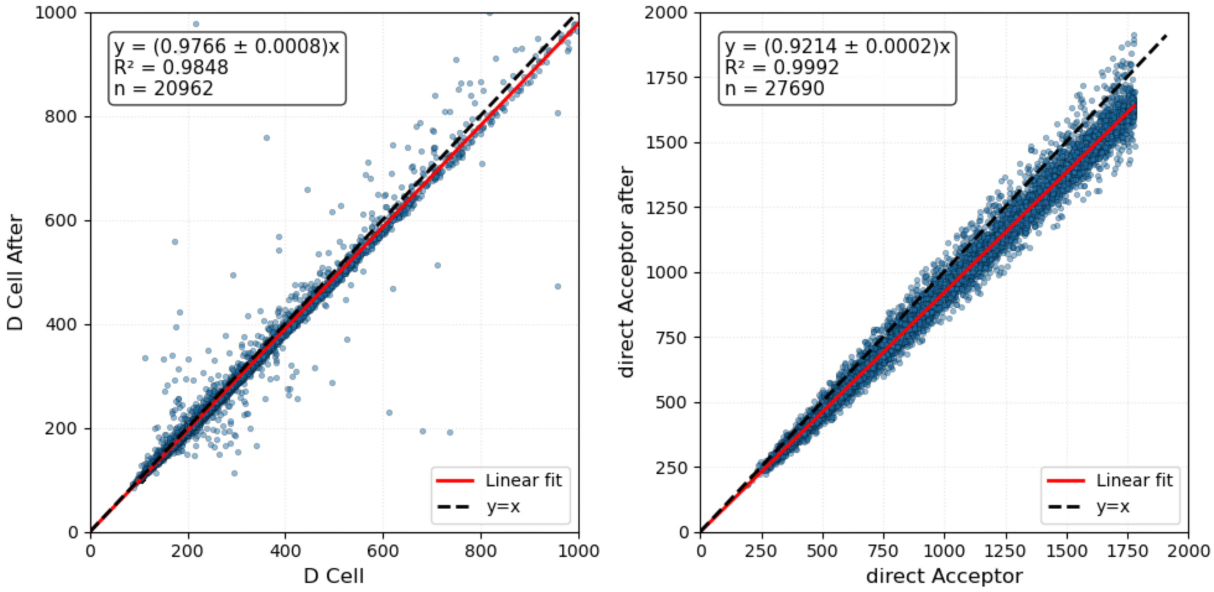


**Fig. S20. Correction of fluorescence intensity changes following osmotic perturbation**

Linear Regressions used to calculate correction factors due to the decrease in fluorescence intensity following osmotic perturbation. The left graph is the donor channel before vs. after for the uncoupled mTurquoise2 fluorophore, while the right graph is the direct acceptor channel before vs. after for all datapoints (cells) of experimental constructs after image analysis filtering (**Figs. S21-S22**). We would expect an average relationship of y = x for both graphs, and we would treat this behavior as an artifact of microscopy. Corrected FRET donor and acceptor after perturbation are calculated from these regressions as:

$$Acceptor_{corr,after} = \frac{Acceptor_{after}}{.9214} - (.070 \times \frac{direct\ Acceptor_{after}}{.9214}) - (0.4163 \times \frac{Donor_{after}}{.9766}),$$

and

$$Donor_{corr,after} = \frac{Donor_{after}}{.9766}.$$

Fluorescence artifact correction factors (**Fig. S19**) remain the same after osmotic perturbation.

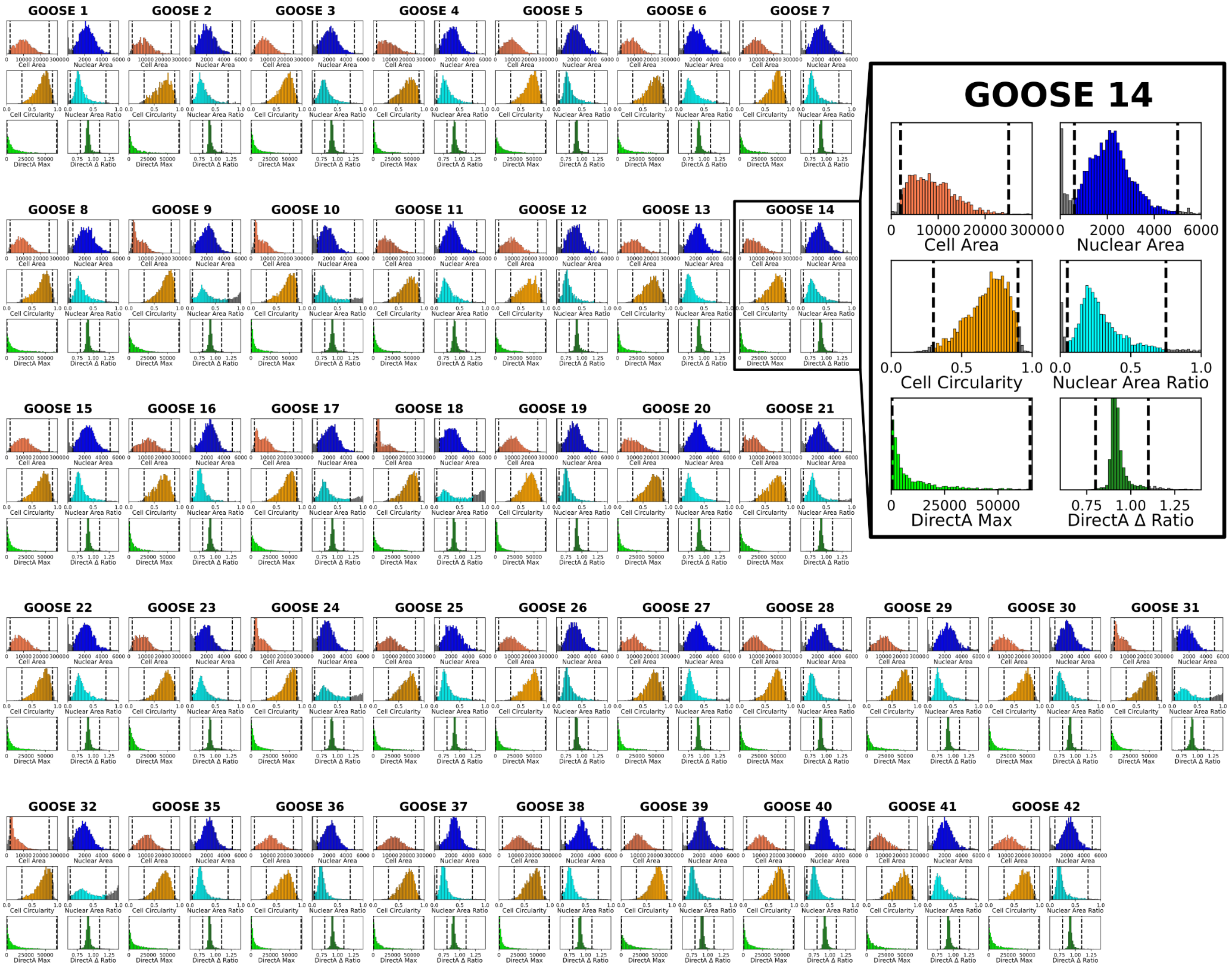


**Fig. S21. Quality control filtering of cell morphology and expression levels**

To characterize the GOOSE-generated constructs, we analyzed the distribution of cell morphology and expression levels. To ensure data integrity, several quality control filters were applied. Cell area was derived from segmentation of the donor channel, while nuclear area was determined via the Hoechst channel. To exclude sub-cellular debris and multi-cell clusters, only cells with an area between 2,000 and 25,000 pixels were included. Similarly, a nuclear area filter (600–5,000 pixels) was applied to ensure a single, well-defined nucleus. To exclude apoptotic or detached cells, circularity, defined as $\frac{4\pi(cell\ area)}{perimeter^2}$, was used as a filter. Cells were required to fall within the circularity range of 0.3 to 0.9; values outside this range typically indicate dying cells or artifacts. The nuclear area ratio was defined as the proportion of total cell area occupied by the nucleus. To filter out cells with abnormal morphology, incorrect segmentation, or minimal cytoplasm (often indicative of cell death), a nuclear area range of 0.05 to 0.75 was used; cells that fell outside this range were discarded. To ensure a robust signal-to-noise ratio and avoid sensor artifacts, cells were filtered based on the maximum direct acceptor intensity (DirectA max). Cells with signals indistinguishable from background or those reaching camera saturation were excluded, using an allowable range of 400 < DirectA max < 65,000. To account for cell loss during the workflow, the ratio of the mean direct acceptor intensity post-perburbation to the pre-perturbation value was calculated (DirectA delta ratio). The acceptable range (0.8 to 1.1) was determined using the population mean (set to .92); values far from the mean indicate cells that have detached. All cutoffs are shown as dashed vertical lines on the distributions.

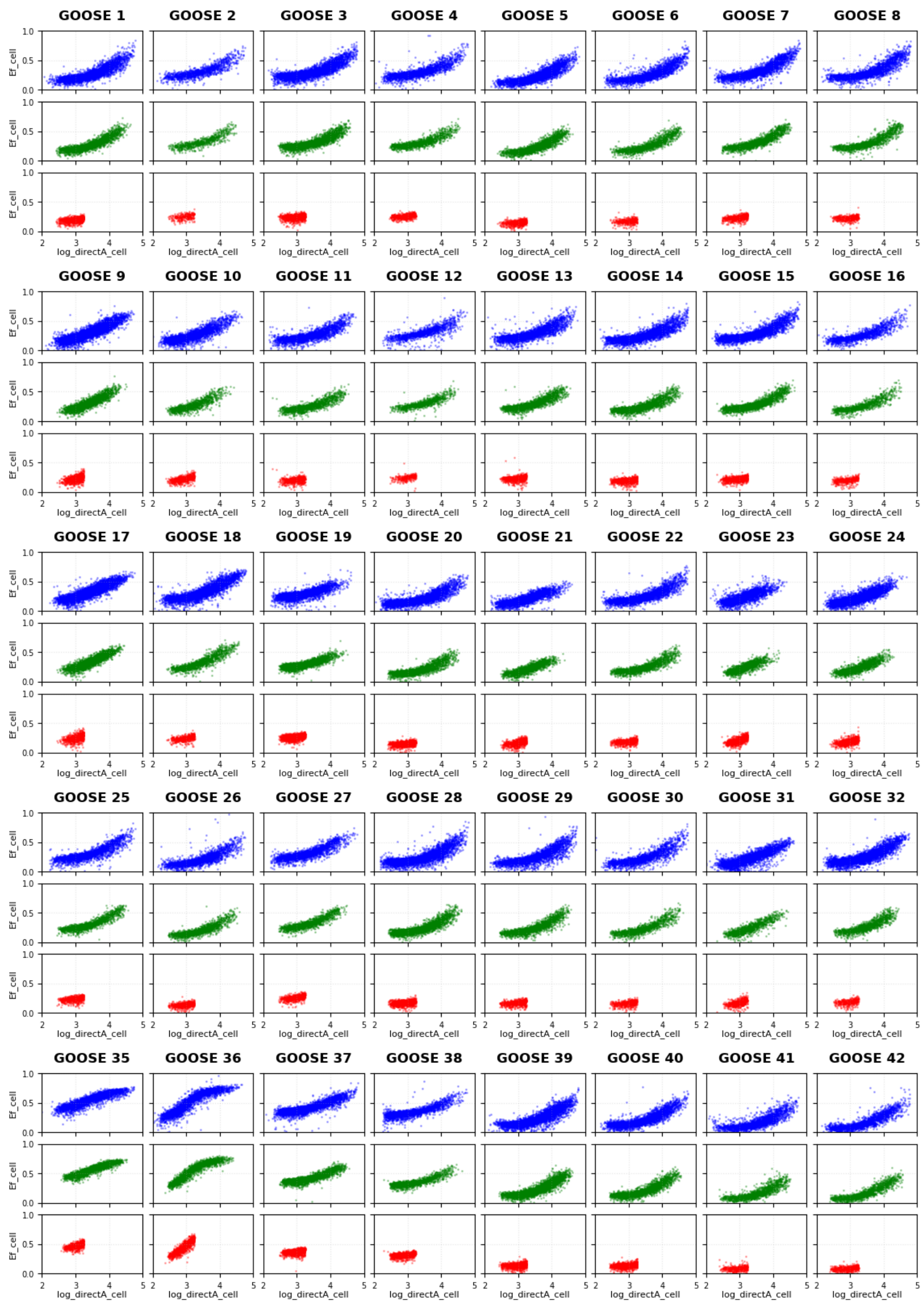


**Fig. S22. Concentration-dependent FRET efficiency and monomeric signal filtering**

FRET efficiency as a function of fluorophore concentration (measured by direct acceptor fluorescence, directA_cell). Each datapoint is a cell. The top row, in blue, is the unfiltered data concatenated from experimental results. The middle row (green) shows the filtered data after removing cells that fall outside the ranges from the previous filtering step (**Fig. S21**). The bottom row (red) contains the finalized data used to calculate violin plots and subsequent average $E_f$ and localization metrics in **Figs. 2, 3**. We remove cells with high fluorophore concentration to ensure we measure monomeric FRET efficiencies 2 < log_directA_cell < 3.25.

| # | Sequence | FCR | NCPR | κ | H | Disorder | Self ε |
|---|---|---|---|---|---|---|---|
| 1 | QNNNQQQNQQNQNNQNNNQNNNNQNQNNQQQQNQQQNNNQQQNNQQNNNNQQQNNNQNNN | 0 | 0 | -1 | 1 | 0.88 | 8.19 |
| 2 | THNHHSTPGTPGHHHPGSPHSPHPTHTTPSHHGTGGGHGGSTTQSHSNGSATGQHGSSGP | 0 | 0 | -1 | 3 | 0.89 | 3.22 |
| 3 | THNHHSTGTGHHHGSHSHTHTTSHHGTGGGHGGSTTQSHSNGSTGQHGSSGPPPPPPPAP | 0 | 0 | -1 | 3 | 0.91 | 3.22 |
| 4 | THNHHGPSTGTGPHHHGSPHSHTHPTTSHHPGTGGGPHGGSTPTQSHSANGSTGPQHGSS | 0 | 0 | -1 | 3 | 0.9 | 3.22 |
| 5 | HNNQQQQQNNQNNQNQQNNQNNNNNQQQQNQQEDEQQDDENQDDDQQDEDQNEDEQQDDE | 0.3 | -0.3 | -1 | 1 | 0.94 | 32.12 |
| 6 | EDDEHNNQDQQQQENNQNDNQNQDQNNQDNNNNNQQQEQNQQDQQNQDQQQNDQQEEDED | 0.3 | -0.3 | -1 | 1 | 0.92 | 32.12 |
| 7 | KKKRSGQNRNQNNRQQQQKNQNQRQNQNKNNQQNQNNENNNNDNNQQENNQQENQDDDDD | 0.3 | 0 | 0.53 | 1 | 0.92 | 8.36 |
| 8 | REKRSGQNKNQNNRQQQQDNQNQEQNQNDNNQQNQNNDNNNNKNNQQDNNQQRNQEKDKD | 0.3 | 0 | 0.13 | 1 | 0.95 | 13.28 |
| 9 | KKRSGKRRHHKKRHQKRRQHRKKQQRKKQNQNNNNNQNQQQQHHNQQNNQNNPNNNNHHP | 0.3 | +0.3 | -1 | 1 | 0.85 | 16.45 |
| 10 | KRKKSGHHKHQQHRQQQNKQNNNRNNQNKQQQQHHNQKQNNQRNNPNRNNNHRHPKKRKR | 0.3 | +0.3 | -1 | 1 | 0.9 | 16.45 |
| 11 | GQSTSTSWGGWGTSGSGGGTSSGGWSSTTSGGEDDGTDDEGTDDETADDDSTDEEGSDED | 0.3 | -0.3 | -1 | 3 | 0.94 | 22.42 |
| 12 | EDDDGQSTESTSWDGGWGDTSGSDGGGTDSSGGWSSTDTSGGEGTGTDTASTDGSEEDED | 0.3 | -0.3 | -1 | 3 | 0.94 | 22.42 |
| 13 | KKKKGHTGRTGTGRGTTGRSSGARGGSARTTTTTSSSSESGSSDSSSAETASSDSSDEEEE | 0.3 | 0 | 0.53 | 3 | 0.95 | 5.68 |
| 14 | KEDKGHTGDTGTGRGTTGRSSGAKGGSAETTTTTSSSSESGSSRSSSAKTASSESSERRED | 0.3 | 0 | 0.13 | 3 | 0.95 | 7.95 |
| 15 | GHTGTGTGGTTGSSGGGSTTTTSSSSSGSSSSSTSSSSDKEKDREAKERDAREREKEAAR | 0.3 | 0 | 0.01 | 3 | 0.89 | 11.23 |
| 16 | GHDTGKTGETGKGTDTGRSSEGGAGSKTTETTRSSDSSASGRSSESSRSTESSKSSEAAR | 0.3 | 0 | 0.06 | 3 | 0.95 | 3.25 |
| 17 | KRRNQKRRGTRKRSGKRKGSRKRGARRKSTATATGSTTSSATASASSSSSSSGAGATGGS | 0.3 | +0.3 | -1 | 3 | 0.94 | 13.60 |
| 18 | KRRKNQGTKSGGSKGASTRATATRGSTTKSSATASASRSSSSRSSGARGATGKGSRRKRR | 0.3 | +0.3 | -1 | 3 | 0.94 | 13.66 |
| 19 | KKRKAHHNKNNQNRQNNNRNQNQRNQNQKNQQQDDEEEDDDDDDDEEDDDDDDDDDDDEE | 0.6 | -0.3 | 0.84 | 1 | 0.95 | 21.22 |
| 20 | DKDAHDKRHNKEDNNKDDQNEDEQNDDDNNNQDEKNQRDDNQERENQDEDNQDDDQQRDD | 0.6 | -0.3 | 0.19 | 1 | 0.97 | 39.66 |
| 21 | KRKSNRRRPPRRKNNKKRPNRRRQQRKRNNPQDDENPEDDNNDDEQQDEEQHEDEPQEEE | 0.6 | 0 | 0.64 | 1 | 0.96 | -11.82 |
| 22 | DDRSNERRPPERENNEKKPNEREQQDKKNNPQKRRNPDDRNNDRKQQEREQHRDEPQDRE | 0.6 | 0 | 0.06 | 1 | 0.96 | 18.49 |

| | | | | | | | |
|---|---|---|---|---|---|---|---|
| 23 | RRRKKKKTKRRKMRKRRPRKKRNKRKRPRRRKPQQNNEQQ NNDNQNNENQNSDQSEDDDD | 0.6 | +0.3 | 0.72 | 1 | 0.94 | 4.03 |
| 24 | DDRTMRRKPNKKEPPRRRQQKKDNNRRKQQNNRRENQRRD NNRKKNQKKRNSKKDQSRDE | 0.6 | +0.3 | 0.22 | 1 | 0.91 | 15.75 |
| 25 | KKKKMVVSRSVVGRAGMARSVVAKSVVARPVSGDDDEEEE EDDDEDDDDDEEDEDDDDDD | 0.6 | -0.3 | 0.84 | 3 | 0.86 | 22.64 |
| 26 | DDKEMVDKDVSEEKSVDEDVGKRDAGDDDMASVVDDASVD EVAPREVSGERDRDKDEDED | 0.6 | -0.3 | 0.16 | 3 | 0.92 | 46.66 |
| 27 | KRRAVKKKVAKRKSSRKKVVKKRGARKKSVSVDEDVPEDD VVDDDGSDDDVSDDEAVDEE | 0.6 | 0 | 0.64 | 3 | 0.88 | -2.21 |
| 28 | EKDAVDEKVADKKSSKDKVVDDRGAKRDSVSVDRDVPDEK VVRDKGSDREVSEKKAVKRD | 0.6 | 0 | 0.06 | 3 | 0.9 | 29.24 |
| 29 | SSGSSGSSEKDAVDEKVADKKKDKVVDDRAKRDVVDRDVP DEKVVRDKDREVEKKAVKRD | 0.6 | 0 | 0.05 | 3 | 0.84 | 29.69 |
| 30 | SESKGDSASVGDSESKVADKKKDKVVDDRAKRDVVDRDVP DEKVVRDKDREVEKKAVKRD | 0.6 | 0 | 0.05 | 3 | 0.86 | 27.54 |
| 31 | KRRKRKRVRKRRVKRKKAKRRRAKRRKVKKKRSAAVGEAV AADSAVADAVVVDSADDDDD | 0.6 | 0.3 | 0.72 | 3 | 0.76 | 9.82 |
| 32 | KRKVVDRDAARRRVSDRKAAKDRVGKRKAVAARKKSARKD VADEKAVRDRVVKRKSADRK | 0.6 | 0.3 | 0.18 | 3 | 0.61 | 28.31 |

**Table S1. Library variants described in Fig. 2 and Fig. 3**

Columns reported here include the Fraction of Charged Residues (FCR), Net Charge Per Residue (NCPR), charge-distribution metric kappa (κ), Kyte-Doolittle hydrophobicity (H), average disorder prediction (Disorder), and a self-interaction (self ε - lower means on average stronger self-interactions). Note that a sequence must have both positive and negative residues for κ to be calculated; otherwise, it is reported as -1. Disorder here is calculated as the average disorder score as predicted by metapredict v2.0. Self-interaction score (self ε) is calculated using the CALVADOS version of the FINCHES model.

| # | Sequence | $R_e$ Target (Å) | Self ε |
|---|---|---|---|
| 35 | EGNDQKQGWTQDNQWWWYNGQWFYPGQFFYQNNGQFQ NFWNFWPNYQQGFYGDNNGFEGE | 31.25 | -16.22 |
| 36 | QPFFGNGNFQTYFQWNGFYGNFKYFGQGYNKWGGGGF NNNFNYGFGNWFETFNGSNGPGQ | 31.25 | -20.91 |
| 37 | FYNWNTNSEDSGHPAGWEFQKPSAKIPELDDTPHEWD DSMSAYSSARNPPWTPALMAYNE | 47.5 | 2.47 |
| 38 | SEENEEPSAEGDWSPDEEFEAGSKTQKYTEDKHSHPD SDKNFSYFPRMKGGSLNSWASSF | 47.5 | 7.75 |
| 39 | TSPPPDDPDSDDTEDDPTTKTEDDTDTTPPTPTSEPP SESESKKEEESPKPTKTPSPTKE | 63.75 | 28.13 |
| 40 | ETEEEEPETPKDESTPTETSSTPTTEEDSEPSPDSES PTKKEPPSKTDSEPKPDPTEEKS | 63.75 | 31.62 |
| 41 | SPDDDDDTDDPETEDDSEDTKESEDSPESDDEEDDDE TPPEEPEDETDESTEDEEEEEPP | 80 | 80.98 |
| 42 | DPTEEDETPEDPDKEDDEDDKDSDTPPPPEDEPPSDP DPEDTDEEPPEDSPEDPEPEPEP | 80 | 69.61 |

**Table S2. $R_e$ designed variants as reported in Fig. 3**

Design objective end-to-end distance ($R_e$) and self-interaction score (ε) calculated using the CALVADOS version of the FINCHES model.

| # | Name | Sequence | Self ε (CALVADOS) | Self ε (Mpipi) | Fluorophore Used |
|---|---|---|---|---|---|
| 43 | Attractive 1 | RWRSYRYRFGRSRKYQQNYWRSNKYFGNGRKGYRSWRRNQYKFQFKY<br>GNNFSWRNGFNQYGRFWSNYQYYGRNYWWNSGRKYNKSRYQWKGRYR<br>YRWRNWYKGQRFQNGSYNGSSWGNNNWGYNWYNWNYWQSYRRKFGQW<br>RKFWNQWWYNQQWQWYKRSQKQNWRKWSGWNGGNGRRYRGWQGKFSR<br>SWYRQYYKQWNY | -74.27 | -30.57 | mTurquoise2 |
| 44 | Attractive 2 | WKFFHWRYYHRWYYFYGGRRKKFRRFGQGGGYSYWRFHYWYWRFGQW<br>QRWGKRFWGKGSWRKRRGYGYGGFFGQKGKGWYRRNSKRFHFRGGFF<br>KFGSRQFRQGRWQGQKGRKWQWRYRYRGRGWSSGRHGFRKGGHRSGY<br>RQGKRYGFRWRWWGRHFYFWYSHFYFFHRWYRRGYSFWKYRGGFKGS<br>RKQFFGSKGYRG | -74.95 | -8.16 | mTurquoise2 |
| 45 | Repulsive 1 | DVDITVIEEEKPTIVKKLPLVKTEVKLAPKVPKPTTIATKTLEELKK<br>DDKVMEPLTIVPDKTDKAITTEVMDKDIPDMEKKLLKKDLMTLMEKE<br>ITTVVPLIKVKTKLVEADTPLTTDTDTAPDIIKDEKKVDDAKDMETD<br>DTKVLLDVILDDTTLLDMLDVVDITDVPKVTTEVDEDMDVPDELMTE<br>LTTTIKLDDLEP | 51.65 | 31.29 | mTurquoise2 |
| 46 | Repulsive 2 | VDVLMTTKVVPLTPELTDDDKEPEVDVVLDADKEDMETEDMKAKTLD<br>LPPIEDVVDKDDLIPKEDDDDVEEDDVTTVETDMPVEEVTAEPTAEP<br>VTDMEKMLKDDTTVIDEDTDIPKVIDADTEDITMEKLKTPEPEDTVV<br>EEELVTADDMDDPTIKMEPMDKTVEKDKVDKDLLKKLAIVKETLPTM<br>PTIDILMDVLAD | 103.45 | 55.08 | mTurquoise2 |

**Table S3. Homotypic designs as reported in Fig. 4d**

Sequence designs in which the target design parameter was self-ε. Sequences 43 and 44 were designed to be self-attractive (negative ε), whereas sequences 45 and 46 were designed to be repulsive (positive ε). Sequences were designed using the Mpipi FINCHES implementation.

| # | Name | Target Scaffold | Sequence | **Self ε (CALVADOS)** | **Client - Scaffold ε (CALVADOS)** | **Self ε (Mpipi)** | **Client - Scaffold ε (Mpipi)** | **Fluorophore Used** |
|---|---|---|---|---|---|---|---|---|
| 47 | Scaffold 1 | N/A | NSQRGNQNYNFWRYNQVYLDTWQIQLFY<br>RRNWVFFFKQNQWPSQIIQNNQAWQNNA<br>LWSFWQNAIQNECQQGQVIMVYQNNIEA<br>QNYRFFNFLILKEQQLQNQLDGRNFYQT<br>WVNFQNEVYVLYYRAEIQLQNNQFQQLM<br>FLCIQMQHQIDSQFNYMMMNHLNMNTYI<br>KENQAVMLGQYQQQYWFIMFYINIAQWV<br>FNYWLYFYWYQMWYNFPNFQLQVFLQNK<br>FNQKLPLGYTQWPNFKQQFQKVAFYQWQ<br>IELNRLMQFQNVILQNCSVWEYYFLFWW<br>QWNAYQFNIMNAVYVLWQVFMMTQYVQQ<br>WYQWHKQWNFRMLYQNIFAIREMWYIYQ<br>QNYLLQVALVQIMWQKQQFVMQQNFYLN<br>IIQFNIMYQPCQFMMMNLAQVWQFVLVL<br>YNYQNRRV | -120.85 | N/A | -70.0 | N/A | mTagBFP2 |
| 48 | Client 1 | Scaffold 1 | FVKAPSFSGGMMCPRFWKFPRYMWPRLL<br>GPRFVSRPLYPPSYAKVRLVANQPPNKR<br>TRRWQSFGFRVRMSRAAPKGRFRVYNWL<br>NSKPYRPRPIRRGHRKLPFALRQMAWQL<br>MKRTGLFIVRRPKRFWHFMRRWFRKNIN<br>LAGQLVMYPPVIPRRPQFFIGFVPKRLW<br>PKRMPLRLKVQRRRGRFFNNLRYRQYKG<br>GRVR | -23.13 | -113.2 | 10.0 | -52.02 | mScarlet3 |
| 49 | Client 2 | Scaffold 1 | RARPFPWAPRGNAWVKPARLQQKQILMS<br>LRLFVRAYQPPCRVGAGSFQVQIFRKVQ<br>FPHVFPRRQPPPYTYTSAGYAKATRFHI<br>GYGPRPGTTPKFCYFPWPQRFMLPPLFW<br>RIAKTRPAKSKVFTVMFKWRWLPVVWRY<br>FRRRGFRYRLPGLRGIYRSYFKGGFRMM<br>QVPRRKAKKFRRMKFGPPVAYRAPPPWL<br>LGPY | -25.21 | -102.94 | 0.04 | -52.04 | mScarlet3 |
| 50 | Client 3 | Scaffold 1 | RGRPFPNTPRASAEGKPPRGGPKNMLSV<br>GRLFSRSYQPPCRGPTSPYPGIPFYKVN<br>YPHGYPRYMPPPYTYTAGFYPKVSYFHS<br>SYIPRPGTMPKFCYFPWPQRYSMPPLFW<br>RGPKTRPPKSKGYTLMYKNRNMPIVERY<br>YYYRMFRYYMPSGRYGYRPYYKSGFRMS<br>QMPYRKGKKFRYMKYGPPGIYRGPPPDG<br>GSPY | -38.53 | -101.22 | -10.05 | -51.95 | mScarlet3 |
| 51 | Not Client 1 | Scaffold 1 | LTGAAETSTASGNTADGDPATGSVGTSD<br>KVSRDVTMSDSITLKSSVEGAKKKSKSS<br>SVGSSAPSITSLASGSSDIKNAMKGGDK<br>DASTTSTTSTKTTVKTSTAKSTKKQTSL<br>TFAKLDTGTDKTGGSVTKIKDAGTLGDS<br>SDTGASSDTKTSTGITTKTKDKATDDPK<br>TSKSSKLDSGKKTTKSTKTSVGQDSGLG<br>AKST | 45.73 | 2.77 | 27.21 | 4.0 | mScarlet3 |

| | | | | | | | | |
|---|---|---|---|---|---|---|---|---|
| 52 | Scaffold 2 | N/A | MPEQNPRFPWFDPYGWGRYRQQPFSEYS<br>GNNNWQPGQRQNQDPWGNQPEQYEQEQE<br>PDNQRQNYQQNWWWQQQPEPQNGNPNQK<br>GNQGQQENFPNNQRGRPRQQYQQNNGQE<br>PNQQINCEQRGRNQRWQRNGDRQNEHPN<br>GERGPHQNWQQNPDGNQQTNQNFNYRGN<br>MQPNQQFDPNNEQNPQPPENFEPNQPRR<br>NQGWWQPQPRQPGPARRNEPNFGQQRNP<br>QQRPRQNWQFYWFRQGGNNYQPNGQQQQ<br>PEFNNDNQDPNNQRQGGRWWNNRQQDPN<br>QRQVDSNQFQNRQNRPRNEFNNQNDQNE<br>GPQSGPRQQGGNRNQPQQPYVRNRLEYE<br>FQQDYNQNNNNRQGEEQRQSQNIRNFQN<br>QRPQGNRQFPGPQPQQERNYPQQYNNPR<br>YPWDYQQQ | -36.5 | N/A | -67.12 | N/A | mScarlet3 |
| 53 | Client 4 | Scaffold 2 | SLINFINNNQLGHENVNTDHGVFNGSSD<br>QMMNHPLPDDGYQQVFPNGMNIMQDHTS<br>AVPGYGPHWGDGVPFGFSNPGHFYNQHD<br>QVPVQLNHPLPFFHVFFVPVPAGWGGQN<br>DQFFFGGWDQVIWLPVQKFQLDILTIDN<br>PPHFEGFLLLDNVNMGNCYQNMNLHGDH<br>FGGNYHMLQSNYPSYLDLYGQHYPPDPF<br>SGNYQQVLYHILQLYENPNDWDFDGNTG<br>DNNDMDHHNWEGSFASQNPFILMQPIGN<br>FVDDGLDQGHNAQQHGNDEDDNHEFNLG<br>NGMVPQNDSLPYNATDGQLPSSSQILNN<br>TMDHHLQFLFLQQLQNNQDVGHRQTQHG<br>NQNFHPPLLNGMHMPQHTALNHSHDEIM<br>QFPWHGFGFPHLPVRQDNQDNLGTYPPV<br>VMQGWHHP | -12.9 | -27.85 | -30.0 | -48.64 | mTagBFP2 |
| 54 | Not Client 2 | Scaffold 2 | GSMGKESNTSGGDAKAIESKMIVLASII<br>MPGDREAADPLPSCVPGMQSPLGAFKAS<br>PTESVQSISSKTDSGDEMKWYTLPDGDS<br>MLDEQDNTPPVMMAEESNLLKTTVIWSK<br>SSNGKMNKMIPTLTASSEPSLLPKPDSK<br>SKDTCMIRVICKSPISTNGPASYSLGMT<br>TTSEDEIAYITFQDSTVGKGEPTIGKSP<br>KSGHNGTVPMLVGTVTVGTGSTSSLLQW<br>NVTPQEMGWKQAEDRNLDDVTTGQNEGM<br>YKDLLPMAKQSEQSEADVGGGNDEADNQ<br>AGMASENSMSMPSQKQVPLGVSSGVTST<br>DSAHIAIMTLVPSQPEMTFTEQTLTPGT<br>ETQDMDSTGEAYSDGNHVQSVSIPSLED<br>VGSEGGEDSHQAGDGTFCLWKNGALESQ<br>IGMSEQIA | 50.72 | 7.64 | 29.95 | -18.06 | mTagBFP2 |

**Table S4. Heterotypic interaction designs, as reported in Fig. 4f,g, and i.**
Sequence designs where the target design parameter was self ε or ε with a target scaffold. Both Scaffold 1 and Scaffold 2 were designed by constraining self ε. Client 1, Client 2, Client 3, and Not Client 1 were all designed by optimizing for ε between themselves and Scaffold 1. Client 4 and Not Client 2 were designed by optimizing for ε between themselves and Scaffold 2.

| # | Uniprot | Sequence | Origin | Organism | References |
|---|---|---|---|---|---|
| 1 | N/A | MLVAIHLLPEKSKMATIVVEKESKS<br>PLADHAPEKSKMARVAIKVTEVHND<br>TGEEQDKKKMM | Synthetic (deleterious) | N/A | |
| 2 | O82355 | MYILKSATRTIASGTIPDPGSLVGS<br>GTTVLDVPVKVAYSIAVSLMKDMCT<br>DWDIDYQLDIGLTFDIPVVGDITIP<br>VSTQGEIKLPSLRDFF | LEA (deleterious) | Arabidopsis thaliana (Mouse-ear cress) | https://doi.org/10.1105/tpc.114.127316<br>LEA_2 |
| 3 | N/A | MLKDKAGSAWNQLKDKAGSAWNQLK<br>DKAGSAWNQLKDKAGSAWNQLKDKA<br>GSAWNQLKDKAGSAWNQLKDKAGSA<br>WNQLKDKAGSAWNQ | LEA (motif only) | Hypsibius exemplaris (Freshwater tardigrade) | https://doi.org/10.7554/eLife.97231.3<br>LEA_4 |
| 4 | N/A | MDATRDKLGEYKDYTADKARETNDS<br>VARKTNETADASRDKLGEYKDYTAD<br>KTRETKDAVAQKASDASEATKNKLG<br>EYKDALARKTRDAKDT | Synthetic LEA | N/A | |
| 5 | P0CU49 | MRPADNWAESQKEKAKAGLKDAQAE<br>VGKVAREVKDKAAGGIEQAKDAVKQ<br>GANDLKRSGSRTFENAKDDIQAKAQ<br>HAKSDLKGAKHQAEGV | LEA | Hypsibius exemplaris (Freshwater tardigrade) | 10.1016/j.molcel.2017.02.018<br>https://dx.doi.org/10.1016/j.molcel.2017.02.018 |
| 6 | A0A438HAK2 | MPRGYSKISDHFNRDRGQYVSKGES<br>LRSPLKKKKNTIRGGTGKNAGEAVT<br>ENVSNVTASARAGLKKTKATLEEKV<br>L | Short LEA | Vitis vinifera (Grape) | LEA_1 inferred by homology |
| 7 | A0A6J3ELF5 | MPTGRKQKGTACLRKLRNSASSSGR<br>NLRNWHRTVRSSAFDVPSDLICGSS<br>TQSEEGAPQEGILEDMPVDPDNEAY<br>EMPSEEGYQDYEPEA | Non-Desiccation Related | Sapajus apella (Brown-capped capuchin) (Cebus apella) | Alpha-synuclein isoform X5 |
| 8 | A0A1U8F7A1 | METAKEKAANIAASAKSGMEKTKAI<br>LEEKAERMSTRDPLKKEMATEKKDD<br>KKTAAELNKREAMDQNATASGICIT<br>S | Short LEA | Capsicum annuum (Capsicum pepper) | LEA_1 |
| 9 | A3AHG5 | MKEKLTVSPAATQEHLGGGEERAVK<br>ERAAEKLGGGEERAVKERAAEKAAS<br>VYFEEKDRLTRERAAERVDKCVEKC<br>VEGCPDATCAHRHGKM | LEA (tile 6) | Oryza sativa subsp. japonica (Rice) | https://doi.org/10.1371/journal.pone.0045117 |
| 10 | A3AHG5 | MKASDASEATKNKLGEYKDALARKT<br>RDAKDTTAQKATEFKDGVKATAQET<br>RDATADTARKAKDATKDTTQTAADK<br>ARETAATHDDATDKGQ | LEA (tile 4) | Oryza sativa subsp. japonica (Rice) | https://doi.org/10.1371/journal.pone.0045117 |
| 11 | A0A4D6M434 | MQAVKEKIVSLNATRKARQEAKAEE<br>RAEKEIAKARMDVAREIRLAKEAEA<br>EMDLHVANTGKKG | Short LEA | Vigna unguiculata (Cowpea) | LEA_1 inferred by homology |

**Table S5. Template sequences for protectant library reported in Fig. 5.** Columns reported here include the uniprot code, if applicable, for each template sequence, the origin (if synthetically generated, if a LEA, and if it was a tile of a longer sequence), the organism it is found in, and references if studied or annotated.

### Table S6. Yeast desiccation survival dataset and sequence features for Fig. 5

The full dataset is available as **Table_S6.csv**. Combined sequencing output from 3 independent biological replicates of the yeast desiccation survival assay. Each row corresponds to a unique variant identified by 'Sequence_ID.'

Differential expression columns: log2FoldChange_{rep} is the $\log_2$ fold change in variant abundance following desiccation stress relative to the unstressed control, as calculated by DESeq2[1] for each replicate independently. 'padj_{rep}' is the Benjamini-Hochberg adjusted p-value for that fold change. 'lfcSE_{rep}' is the standard error of the $\log_2$ fold change estimate. These per-replicate values were subsequently averaged across all three replicates and tested for statistical significance to generate weighted average performance metrics. 'Base_origin' denotes the template sequence from which each variant was derived, used to group variants for within-template analyses.

NARDINI+ feature columns (prefix: nardini_): Physicochemical sequence features computed using NARDINI+ and normalized against the S. cerevisiae IDRome[2]. The suffix _raw denotes the absolute calculated value, while _zscore denotes the value normalized as a z-score relative to the distribution of that feature across the S. cerevisiae IDRome, reflecting how unusual a given sequence is relative to the yeast disordered proteome. Frac {X}_raw/zscore is the fraction of residue type X (single-letter code) in the sequence. 'Frac Polar,' 'Frac Aliphatic,' and 'Frac Aromatic' are the fractions of residues belonging to those physicochemical classes. 'Frac Chain Expanding' is the fraction of chain-expanding residues (K, R, D, E, P). 'FCR' is the fraction of charged residues (K + R + D + E). 'NCPR' is the net charge per residue, defined as ($f^+ - f^-$). 'Hydrophobicity' is the mean Kyte-Doolittle hydrophobicity, rescaled to a 0–1 range. 'Disorder Promoting' is the fraction of residues classified as disorder-promoting (A, R, G, Q, S, E, K, P, D, H, T). 'Iso point' is the theoretical isoelectric point. 'PPII' is the mean polyproline II (PPII) helix propensity. '{X} Patch' (raw and z-score) is the patchiness score for residue type X, quantifying the degree to which that residue type is locally clustered versus uniformly distributed along the sequence. 'RG Frac' is the combined fraction of arginine and glycine residues.

CIDER feature columns (prefix: cider_): Sequence parameters computed using CIDER[3]. 'kappa' ($\kappa$) quantifies the linear patterning of oppositely charged residues relative to one another, ranging from 0 (well-mixed) to 1 (fully segregated into blocks of like charge). 'omega' ($\Omega$) quantifies the patterning of chain-expanding residues (K, R, D, E, P) relative to all other residues, ranging from 0 (well-mixed) to 1 (clustered). 'delta' ($\delta$) is the raw, unnormalized measure of local charge asymmetry from which $\kappa$ is derived ($\kappa = \delta / \delta\_max$); unlike $\kappa$, $\delta$ is not normalized by composition. 'length' is the sequence length in residues. 'mean_net_charge' is the mean absolute net charge per residue. 'uversky_hydropathy' is the mean hydropathy value used in the Uversky charge-hydropathy plot for disorder classification. Fraction {x} charge classes were also calculated: negative, positive, and neutral.

SPARROW feature columns (prefix: sparrow_): Sequence parameters and predicted conformational properties computed using SPARROW[4]. 'SCD' is the sequence charge decoration parameter, which captures the linear patterning of charged residues in a manner similar to $\kappa$ but using a different mathematical formalism. 'SHD' is the sequence hydropathy decoration parameter, capturing the linear patterning of hydrophobic residues. 'complexity' is a measure of sequence compositional complexity[5]. Sparrow also calculated fraction {x} physiochemical classes, like positive, negative, polar, aliphatic, aromatic and proline. The 'scaled_rg' and 'scaled_re' are the predicted radius of gyration and end-to-end distance, respectively, scaled by sequence length, as predicted by ALBATROSS. The 'prefactor' and 'scaling_exponent' are the prefactor and Flory scaling exponent ($v$) derived from the predicted dimensions, where $v = 0.5$ indicates a Gaussian chain, $v = 0.6$ an expanded coil, and $v < 0.5$ a compact globule. The 'asphericity' is the predicted asphericity of the conformational ensemble, where 0 indicates a perfectly spherical ensemble and 1 a fully rod-like conformation. The 'avg_helix_prob,' 'avg_beta_prob,' and 'avg_coil_prob' are the mean per-residue probabilities of α-helix, β-strand, and coil conformations, respectively, averaged across the sequence. The 'avg_mito_targeting,' 'avg_nes,' and 'avg_nis' are the mean predicted mitochondrial targeting, nuclear export signal, and nuclear import signal scores, respectively.

FINCHES feature columns (prefix: finches_): The 'mpipi_epsilon_self' is the predicted homotypic interaction energy calculated using the FINCHES implementation of Mpipi (Mpipi-GG) force field, reflecting the propensity for self-association[6,7]. Similarly, 'calvados_epsilon_self' is the analogous homotypic interaction energy, estimated using the CALVADOS coarse-grained force field, as implemented in FINCHES[6,8].


1. Love, M., Anders, S. & Huber, W. Differential analysis of count data – the DESeq2 package. *Genome Biol.* (2013). at <https://cdimage.debian.org/mirror/bioconductor.org/packages/3.3/bioc/vignettes/DESeq2/inst/doc/DESeq2.pdf>
2. Ruff, K. M., King, M. R., Ying, A. W., Liu, V., Pant, A., Lieberman, W. E., Shinn, M. K., Su, X., Kadoch, C. & Pappu, R. V. Molecular grammars of predicted intrinsically disordered regions that span the human proteome. *Cell* **0,** (2025).
3. Holehouse, A. S., Das, R. K., Ahad, J. N., Richardson, M. O. G. & Pappu, R. V. CIDER: Resources to analyze sequence-ensemble relationships of intrinsically disordered proteins. *Biophys. J.* **112,** 16–21 (2017).
4. Lotthammer, J. M., Ginell, G. M., Griffith, D., Emenecker, R. J. & Holehouse, A. S. Direct prediction of intrinsically disordered protein conformational properties from sequence. *Nat. Methods* **21,** 465–476 (2024).
5. Wootton, J. C. & Federhen, S. Statistics of local complexity in amino acid sequences and sequence databases. *Comput. Chem.* **17,** 149–163 (1993).
6. Ginell, G. M., Emenecker, R. J., Lotthammer, J. M., Keeley, A. T., Plassmeyer, S. P., Razo, N., Usher, E. T., Pelham, J. F. & Holehouse, A. S. Sequence-based prediction of intermolecular interactions driven by disordered regions. *Science* **388,** eadq8381 (2025).
7. Joseph, J. A., Reinhardt, A., Aguirre, A., Chew, P. Y., Russell, K. O., Espinosa, J. R., Garaizar, A. & Collepardo-Guevara, R. Physics-driven coarse-grained model for biomolecular phase separation with near-quantitative accuracy. *Nat. Comput. Sci.* **1,** 732–743 (2021).
8. Tesei, G. & Lindorff-Larsen, K. Improved predictions of phase behaviour of intrinsically disordered proteins by tuning the interaction range. *Open Res. Eur.* **2,** 94 (2022).
